# SAMD1 Distribution Patterns in Mouse Atherosclerosis Models Suggest Roles in LDL Retention, Antigen Presentation, and Cell Phenotype Modulation

**DOI:** 10.1101/2021.09.12.459413

**Authors:** Bruce Campbell, Patricia Bourassa, Robert Aiello

## Abstract

The theory that lesions formed by retention of circulating LDL can then progress to complicated atherosclerotic lesions has been a subject of debate, as has the mechanism of retention. In earlier work, we identified SAMD1, a protein expressed by intimal smooth muscle cells in human lesions that appears to irreversibly bind apoB-Lps in extracellular matrix near the lumen. We hypothesized this binding could contribute to the formation of lesions in mice, and that inhibiting binding could reduce lesion growth. In mouse models of atherosclerosis, we found that SAMD1 binds LDL; that SAMD1/apoB complex is ingested by intimal cells; and that recognizable epitopes of the SAMD1/apoB complex survive some degree of catabolism in foam cell. These data appear to support the SAMD1/LDL retention hypothesis of lesion growth. Despite apparently irreversible binding of human LDL to full-length human SAMD1, efficient anti-SAMD1-antibody inhibitors were created. In vivo lesion targeting of inhibitors was demonstrated by MRI, ultrasound, and *ex vivo* microscopy. However, only non-statistically significant reductions in spontaneous lesion size in apoE-/- mice were seen after 12 weeks of treatment with PEG-fab inhibitors of SAMD1/LDL binding. In contrast, these inhibitors substantially reduced LDL retention in carotid injury lesions in apoE-/- and LDLR-/- mice 7 days after injury. The most obvious difference between injury lesions and early spontaneous lesions is the presence of numerous smooth muscle cells and associated extracellular matrix in the injury lesions. Thus, SAMD1 may be involved in retention of apoB-Lps in mouse lesions, but not until smooth muscle cells have entered the intima. In addition, SAMD1 is seen throughout arteries in changing patterns that suggest broader and more complicated roles in atherosclerosis.

## Introduction

The canonical view of lesion initiation is based on the “The Response-to-Retention Hypothesis of Early Atherogenesis” (R2R) [1], in which circulating apoB-Lps are retained sub-endothelially, where they are modified such that monocytes are recruited into the intima, monocytes convert to macrophages, and ingest the retained lipid. Uptake of modified lipids by macrophages can be unregulated and, with sufficient lipid availability, macrophages can become foam cells. The R2R theory has been extended as follows: With insufficient cholesterol efflux, foam cells become apoptotic and release lipid to form lipid pools near the media. Over time, the process continues with additional lipids and cellular debris accumulating to become the necrotic cores of lesions. Most data supporting the R2R theory, and its extension, come from mouse models of atherosclerosis that provide rapid ways to explore and modify lesion initiation and development. The most frequently used are apoE-/- and LDLR-/-mice, which were created in 1992 and 1993 respectively. In these models, genetic modifications, typically with the addition of high fat diets, cause very high serum LDL levels that induce spontaneous aortic lesions. Greatly accelerated lesion growth in knockout models can be forced by various forms of carotid injury, but the lesions are morphologically different from spontaneous lesions.

Previously we described several interesting characteristics of SAMD1 [2]. The protein was isolated from rabbit injury model lesions, and bound LDL *in vitro*. In human lesions, extracellular SAMD1 was consistently colocalized with apoB in intimal extracellular matrix (ECM) around SMCs. Further, recognizable epitopes of SAMD1 and apoB remained colocalized around the lipid droplets within foam cells, suggesting that SAMD1 was involved in athero-related retention and ingestion of apoB-Lps. Colocalization in lesions appeared to be in accordance with the response to retention hypotheses. We thus hypothesized that SAMD1 could be involved in LDL retention leading to atherogenesis and lesion growth, and here report on tests of that hypothesis in mouse models of atherosclerosis.

A recent paper strongly supports SAMD1’s participation in atherosclerosis. SAMD1-miR-378c controls VSMC dedifferentiation, proliferation, and SAMD1 expression, and also demonstrates that SAMD1/LDL binding leads to LDL oxidation and foam cell development [3]. Otherwise, SAMD1 has not been widely studied [4], and reports have primarily focused on roles in epigenetics:

- SAMD1 is a chromatin regulator that modulates the function of active CGIs in mouse embryonic stem (ES) cells for normal embryonic stem cell differentiation [5].
- In a study of regulatory regions that can determine cell identity, SAMD1 was the strongest enhancer in H3K4me3 [6];
- SAMD1’s functional associations and predicted biological processes (GO) with the highest Z-score mostly involve epigenetics, and control of replication and proliferation [7], [8].
- SAMD1 was among the highest 5% of proteins for association with both TAF3 recognized H3K4me3-modified, and bivalently H3K4me3/H3K27me3-modified chromatin. TAF3 is a regulator of p53 which modulates cell cycle arrest and programmed cell death [9];
- Oscillatory cAMP signaling rapidly alters H3K4 methylation; SAMD1 is listed as a gene in which promoter peaks were rapidly down-regulated by cAMP and then returned to baseline quickly after cAMP washout [10].
- It can be inferred that SAMD1 is a methyl-lysine reader which recognizes methylated lysine residues on histone protein tails, and are associated with repression of gene expression [11];
- SAMD1 may be involved in myogenic and adipogenic cell fate commitment [12], and changing cell transcription response to external stimuli [10].
- SAMD1’s participation in epigenetic regulation may be related to changes in cell phenotypes, and is reported to be a participant in H3K4-methylation factors. H3K4me3 is a permanent epigenetic marker of SMC lineage in the SMMHC promoter [13].

## Results

### SAMD1/LDL Binding and Inhibitors

SAMD1 was expressed in HEK293 cells (Fig. 1a). Full-length human SAMD1 was highly expressed, but was difficult to extract, poorly soluble, and possibly bound to cell cytoskeleton; the purified protein tended to aggregate and precipitate unless stored at dry ice temperatures. HEK293 cells would only express mouse SAMD1 fragments; we used the 35kDa fragment (from the N-terminus (5’) start to aa279) in subsequent work. E coli were able to express a human C-terminus fragment that started at aa344 and went to the end at aa538, and e coli were able to express a rabbit SAMD1 fragment from a.a. 238 to the C-terminus (3’) end at a.a. 550 [2].

**Figure 1:**
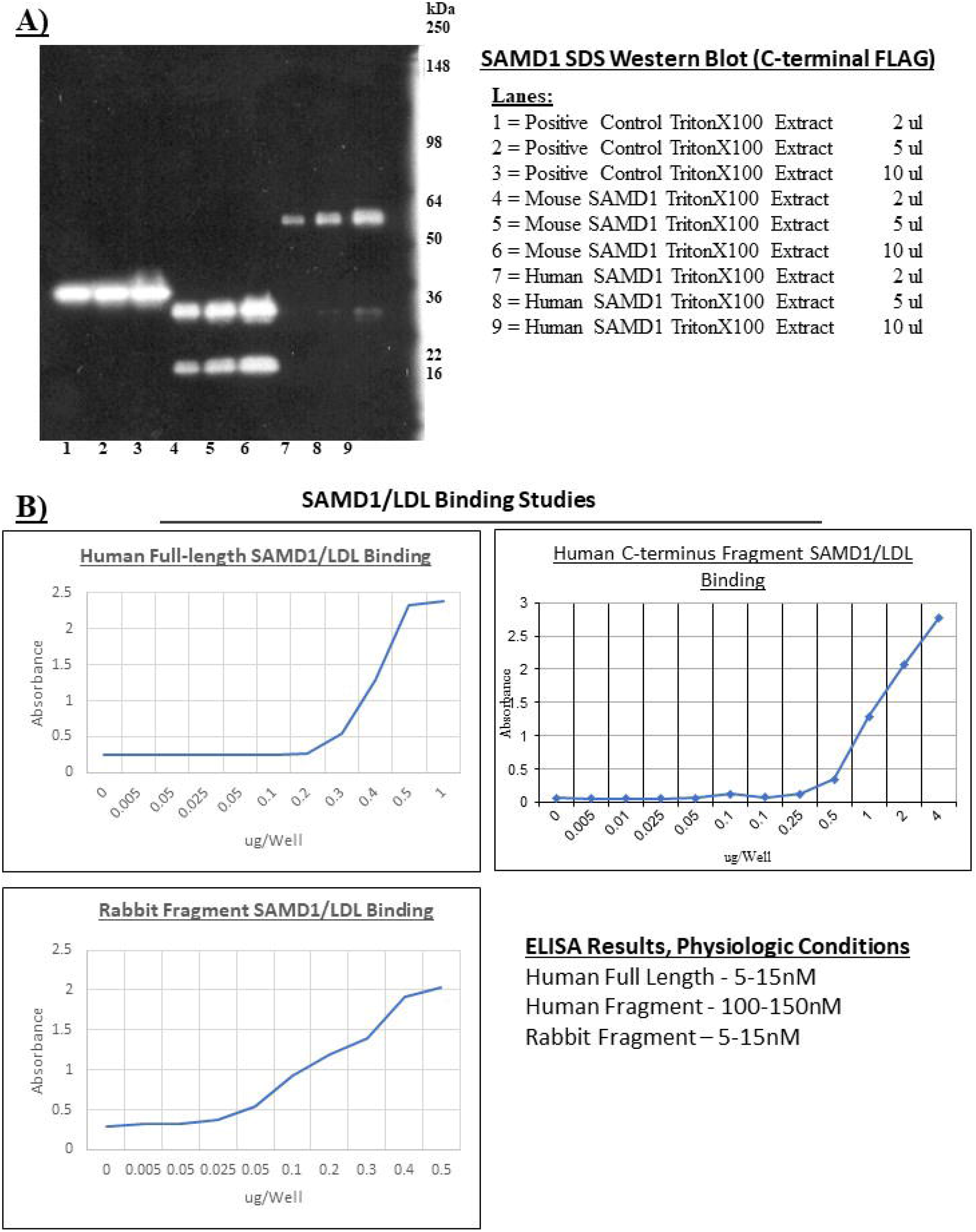
SAMD1 Production and LDL Binding. A) SAMD1 expression in HEK293 cells shows a single 56kDa full length human SAMD1 band, and two bands of mouse SAMD1. B) SAMD1 and LDL bind tightly under physiologic conditions. Freshly isolated human LDL was applied to 96-well ELISA plates and increasing concentrations of SAMD1 were added. By 0.5 micrograms per well, (100-150 nM) full-length human SAMD1 binding to LDL was saturated, and the shape of the curve suggests a Kd of 5-15 nano-molar. A 196aa human SAMD1 C-terminus fragment expressed by e coli has an order of magnitude weaker binding. LDL binding for the 312aa rabbit C-terminus fragment was estimated to be in the 15-20Kd range.

Full-length human SAMD1 bound to freshly isolated human LDL with a Kd of 5-15 nano-molar, as calculated from ELISAs run under physiologic conditions, while the human fragment bound LDL less strongly (Fig. 1b). The rabbit fragment bound LDL with a Kd of about 15nM, but the binding curve suggested somewhat binding different behavior. However, simple ELISAs rely on kinetics models that do not differentiate highly efficient behavior. Interestingly, SAMD1/LDL binding kinetics could not be determined by Biacore, because, although the sensor chip reached steady state during the association phase, dissociation was not observed during buffer flow. This was true both when LDL was bound to the sensor and SAMD1 was the reactant, and when SAMD1 was bound to the sensor and LDL was the reactant; these results suggested irreversible binding. Irreversible binding *in vivo* is also implied by the persistence of recognizable epitopes of SAMD1 colocated with apoB after incomplete lysis in foam cells (Figs. 9,10). Note that SAMD1 did not bind to LDL purchased from four different vendors, only freshly isolated human LDL was functional.

Antibodies (abs) to human and mouse SAMD1 were made in mice, and fab-fragments (fabs) to human and mouse SAMD1 were made by phage display. Seven mAbs and 2 fabs demonstrated varying degrees of inhibition of SAMD1/LDL binding as measured by competition ELISA under physiologic conditions. fab1621 and fab1961, made against full-length human SAMD1, and mAb1A11 and mAb4B6, made against the mouse SAMD1fragment, had low nanomolar IC50s. (Fig. 2a). SAMD1 and inhibitors had to be mixed prior to being added to plate-bound LDL, because inhibitors were minimally effective at displacing SAMD1 from pre-formed SAMD1/LDL complex (Fig. 2c); this result supports the hypothesis of SAMD1/LDL irreversible binding. PEGylation improved the IC50 of both fabs to sub-nanomolar, as well as extending their serum half-life to 17 hours. The truncated SAMD1 fragments were tested with inhibitors in competition ELISAs to narrow the location of SAMD1’s apoB binding; none of the highly effective inhibitors bound to the human C-terminus fragment, but did bind to the mouse fragment and the rabbit fragment (data not shown). ELISAs were used to select inhibitors with strong affinities for full-length human SAMD1; fab1621, fab1961, and mAb4B6 displayed low nanomolar affinities (Fig. 2c).

**Figure 2:**
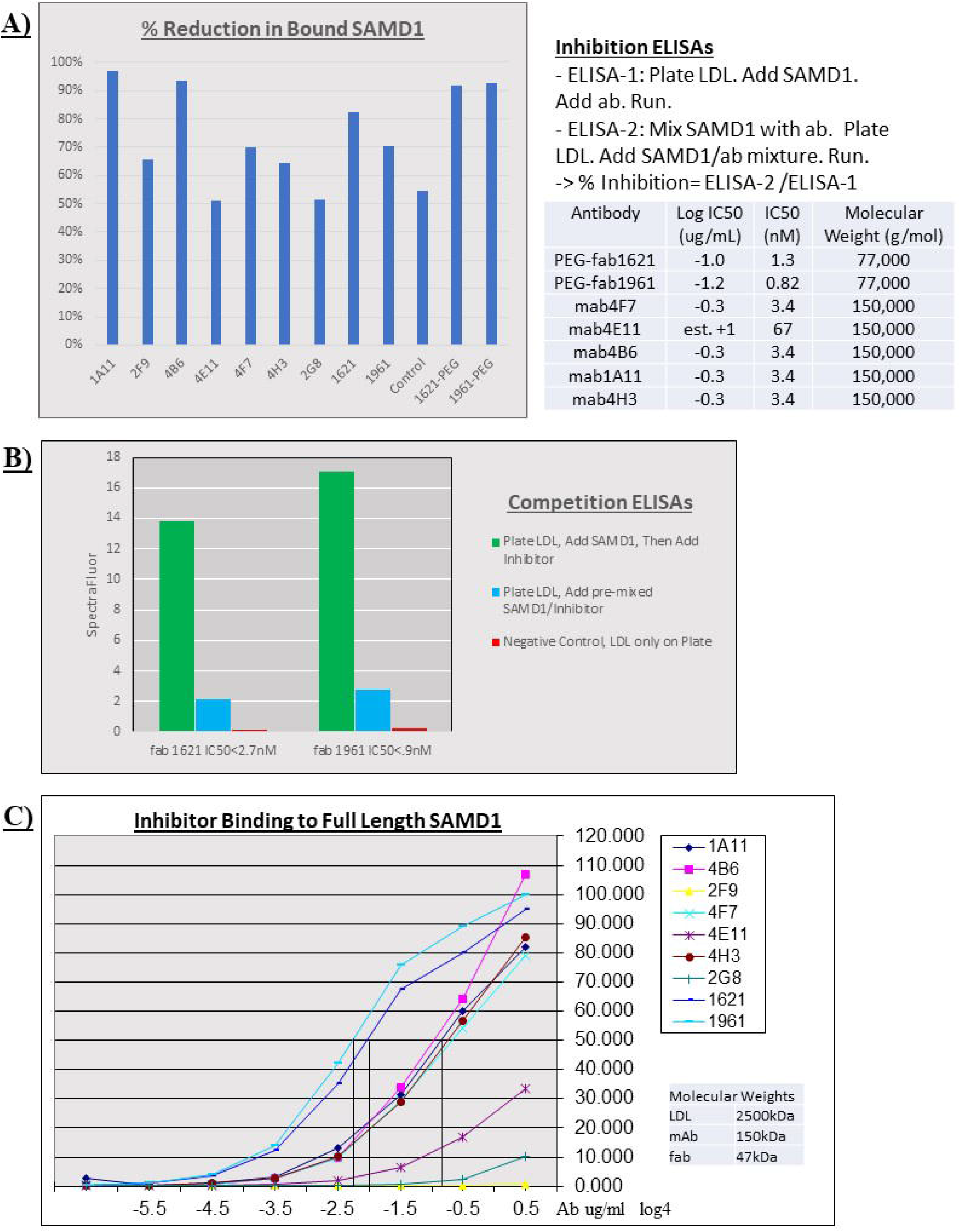
In Vitro Inhibition of SAMD1/LDL Binding. A) Inhibition of SAMD1/LDL binding is shown for each of the inhibitors. Two mAbs outperformed the fabs fragments in inhibition, pegylation improved the inhibition of the fabs, and the fabs had substantially better binding to SAMD1. Although the control, an irrelevant fab, appears to show inhibition, it is actually just showing differences in background signal, because the fab did not bind to SAMD1. B) Competitive binding ELISAs for the 2 fabs shows that inhibitors with single digit nM Kd IC50 do not release LDL when SAMD1 is pre-bound to LDL. The first ELISA used two blocks of fabs in a sequential format wherein SAMD1 was allowed to bind to LDL on the plate first, before washing and adding fab, resulting in a maximum signal that was dependent on antibody concentration and affinity for SAMD1; i.e., no inhibition was observed. The second ELISA was performed by mixing fabs and SAMD1 before adding the mixture to LDL on the plate. The final concentration of SAMD1 is the same in both sequences, 500 ng/ml, allowing comparison between the assays. C) ELISA binding curves for a panel of 7 mAbs and 3 fabs show low nanomolar binding coefficients for the best mAbs (4B6= 3.4nM Kd) and fabs (1961= .7nM Kd), and weak binding for three mAbs.

Performance of the seven inhibitors was compared by IHC in slices from seven different descending artery sections taken from WT, apoE-/-, and LDLR-/- mice after varying times on HF diet (data not shown). Polyclonal guinea pig antibodies against the rabbit SAMD1 fragment (GP-abs) were used in IHC for comparisons with the mAbs and fabs. The mAbs behaved similarly to each other, with stain differing in intensity, rather than differences in distribution patterns, but under-stained in the media and ECs, and stained more strongly in acellular lipid accumulations, when compared to the fabs and GP-abs. Overall, fab1961 stained more strongly than the other inhibitors, particularly in medial and adipose cells. Note that IHC and IF staining for SAMD1 appeared to work with formalin fixation, but not frozen fixation.

### SAMD1 Expression in Cell Culture

RT-PCR showed that gentle abrasion of human ECs in layered coculture above SMCs stimulated upregulation of SAMD1 by a factor of 8 (Fig. 3a). In a comparison of multiple stressors, SMCs incubated with cell culture media from injured ECs may have substantially increased SAMD1 mRNA; but the wide variation between repeat qRT-PCR runs for base SAMD1 makes the result inconclusive (Fig. 3b). Repeated trials added comparisons of different SAMD1 primers, but similarly wide variations between runs were observed (Supplementary Fig. 1).

**Figure 3:**
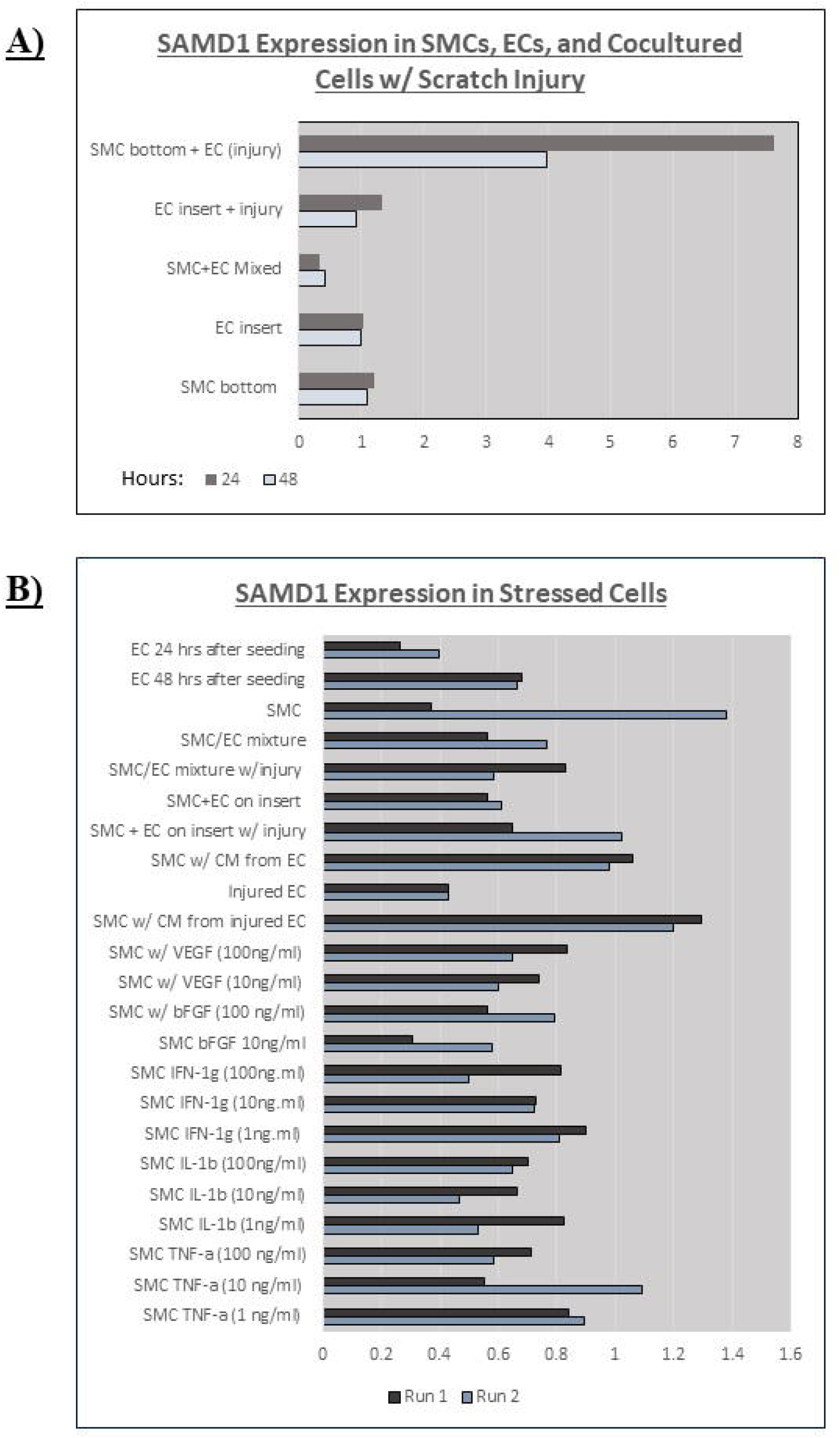
Stressing cultured Cells Changes SAMD1 mRNA Expression. A) Physical stress upregulated SAMD1 expression in layered cell coculture. Human Aortic Endothelial Cells (ECs), and Human Aortic Smooth Muscle Cells (SMCs), were grown in monocultures, in mixed cell cultures, and with SMCs grown on the plate and ECs grown on Transwell permeable inserts. Gentle abrasion caused substantial upregulation of SAMD1 at 24 only when applied to ECs in the permeable inserts above SMCs; by 48 hours, levels had fallen by half from the peak at 24 hours. There was minimal effect on SAMD1 expression in cell monocultures, and in the other coculture arrangements. B) Cultured ECs, SMCs, and EC/SMC cocultures were scratch stressed and mRNA levels were compared to uninjured cells, cells with cell medium from injured cells added, and SMCs stressed by adding increasing amounts of VEGF, bFGF, IFN, IL-1b, or TNF-a to their culture media. Relative SAMD1 expression was measured by qRT-PCR. The experiment was repeated twice, with a very large difference in expression in the control SMCs [Supplementary Fig. 1]. A version of this experiment was repeated 4 times using two different SAMD1 primers, and A) compared to mRNA levels in ECs, whole aorta, and T-cells. In each run, the cytokines had different effects on SAMD1 expression levels (Supplementary Fig. 1)

SMCs were induced to express SAMD1 and secrete it to the ECM, as shown by confocal microscopy using fab1961 as a probe (Fig. 4a). A morphologically different Mouse Aortic SMC (MASMC) towards the edge of the plate expressed SAMD1 and aSMA, which is a marker of SMC differentiation [14]. aSMA also appeared in the ECM of this cell, demonstrating that SAMD1 translocates to the ECM of at least one SMC phenotype, and that SAMD1 expression can correlate with phenotypic differences/changes *in vitro*. Cultured Mouse Heart ECs expressed a small round dot of SAMD1 at the edge of some nuclei, and more broadly in the cytoplasm of some cells (Fig. 4b).

**Figure 4:**
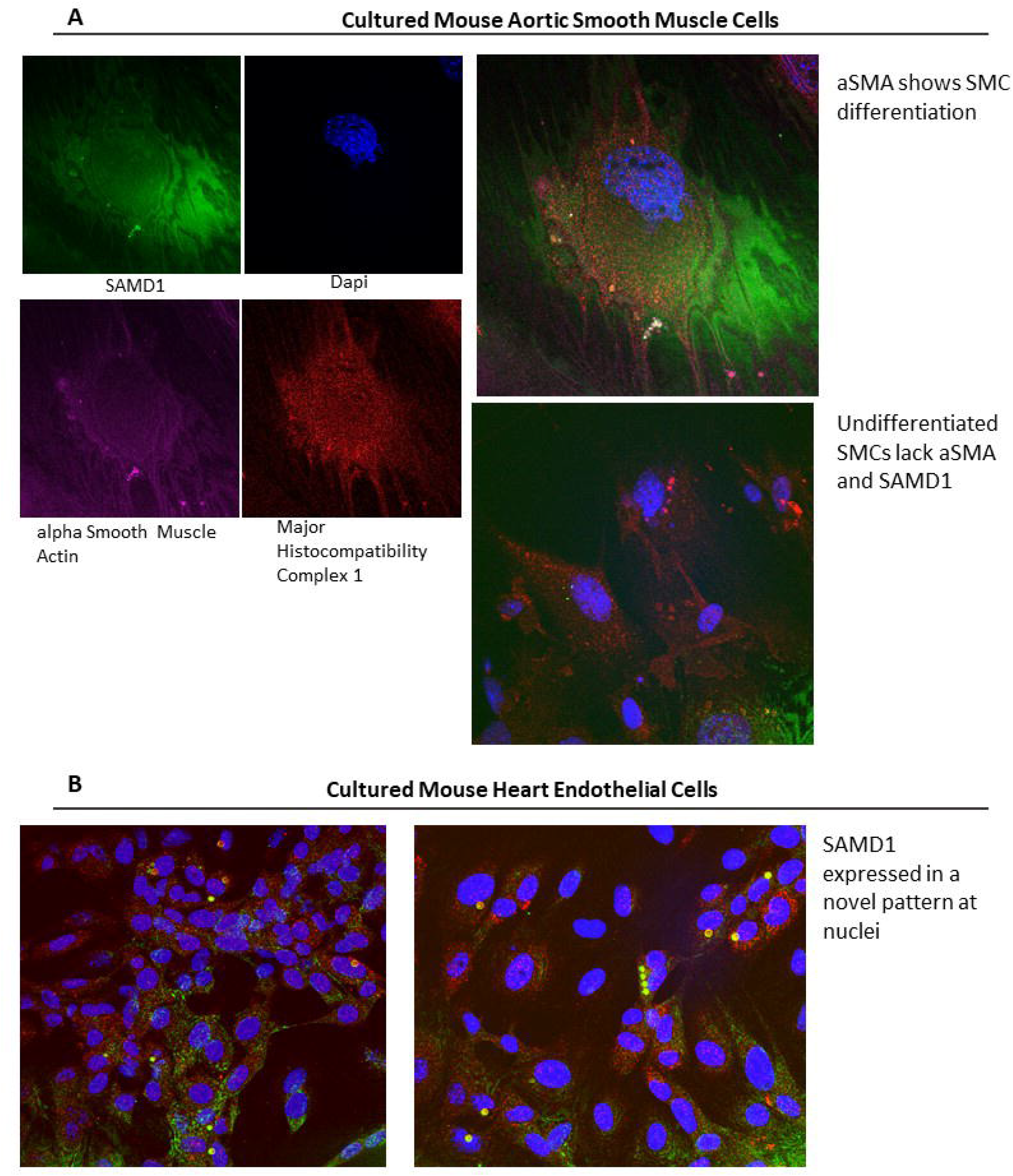
SAMD1 Protein Localization in Cultured SMCs and ECs. Cells were co-stained for SAMD1 (green), Major Histocompatibility Complex 1 (red), DAPI (blue), and, for the SMCs, alpha smooth muscle actin staining was added (purple). A) A differentiated cultured mouse aortic SMC (MASMC) stained for SAMD1 intracellularly and in the ECM. MASMC differentiation is shown by actin expression, MHCI roughly marks the extent of the cell, and SAMD1 is seen intra- and extracellularly. In some areas, SAMD1 and actin appear colocalized, in other locations, SAMD1 is on the surface of the actin bands, as can be seen when the actin channel is intensified. Undifferentiated MASMCs from the same plate lack actin and SAMD1. B) Mouse Heart ECs (MHECs) also stain for SAMD1. On one part of the plate, MHECs appear to show increased SAMD1 expression where MHECs are interacting with each other. Another area of the plate appears to show SAMD1 strongly expressed and colocalized with MHCI in small circles at nuclei.

### SAMD1 Biodistribution

SAMD1 mRNA in WT and LOX-1 mice was widespread, with spleen, blood, muscle, and heart having the highest expression levels, and lower levels in the liver, lung, fat, uterus, aorta, kidney, and macrophages (Fig. 5a). Confocal images showed biodistribution of antibody-accessible SAMD1 protein after *in vivo* injection of Alexa488-labeled antibodies to SAMD1 (Fig. 5b). Fabs were localized primarily in kidneys, liver, spleen, and capillaries of the heart. Subclavian adipose staining by confocal was below levels of collagen autofluorescence, but in several atherosclerotic mice, anti-SAMD1 antibodies also stained perivascular adventitial tissue (PVAT). Measurable SAMD1 protein was not found in mouse plasma.

**Figure 5:**
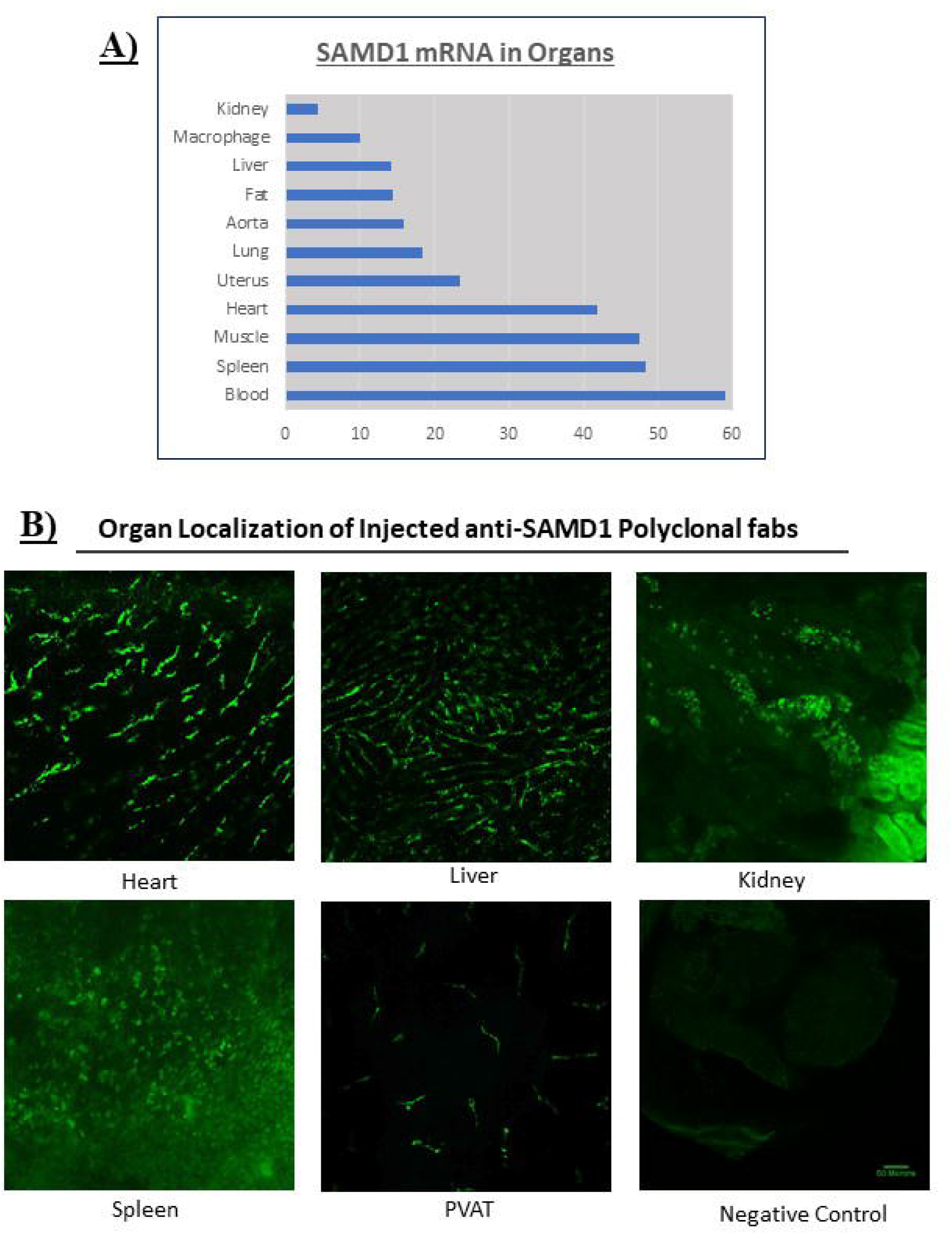
SAMD1 mRNA Biodistribution. A) Biodistribution of SAMD1 mRNA by qPCR. Data from c57bl6 and LOX-1 mice was combined. B) Localization of SAMD1 in organs of 1 year old LDLR-/- mouse. Confocal images taken 24 hours after injection of Alexa488-labeled polyclonal anti-SAMD1 fab fragments shows SAMD1 in kidney, heart, liver, PVAT, and spleen. IHC with polyclonal anti-SAMD1 antibodies strongly stained thymus fragments that remained attached to an aorta.

### SAMD1 mRNA in Aorta

After various times on HFD, SAMD1 mRNA levels were determined from whole artery preparations taken from WT aortas; apoE-/- aortas; and injured and non-injured apoE-/- carotids (Fig. 6). After 4 weeks on high fat diet, SAMD1 mRNA levels in the apoE-/- mice were more than 2X the 10-week WT, but by 10 weeks on HFD, the lesioned and non-lesioned apoE-/- mRNA levels had dropped to almost the same as the 10-week WT, but rose slightly at 20 weeks.

**Figure 6.**
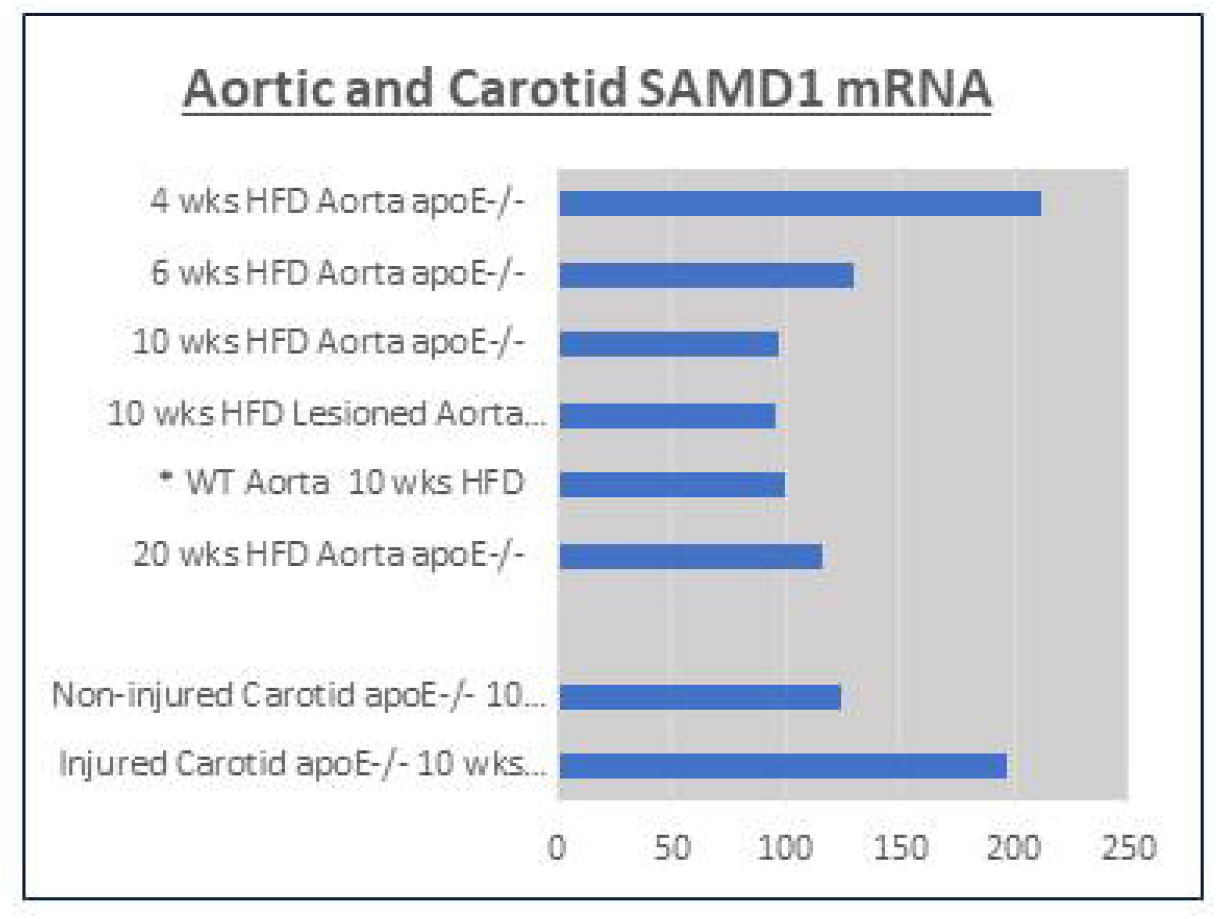
SAMD1 mRNA in Mouse Aortas. SAMD1 mRNA levels were measured in the entire aortas and carotids and were normalized to the WT aorta at 10 weeks to provide a comparison to a stable base state. Amounts of SAMD1 protein in each layer of the aorta varied as lesions developed.

A different response was seen in apoE-/- carotid injury lesions at four days after ligation; here, SAMD1 mRNA was almost 70% above that of the uninjured carotid. Qualitatively, SAMD1 IHC staining in knockout aortas compared to WT aortas suggests that increased SAMD1 mRNA levels in spontaneous lesion intimae may be somewhat obscured by corresponding large decreases in medial SAMD1 mRNA.

SAMD1 aortic mRNA levels were also studied in a transplant lesion regression model, and compared to SAMD1 mRNA levels in spontaneous lesions using non-lesioned transplants as controls (N=5 for each group). Lesion volume regression was quantified; there was no clear correlation between mRNA levels and lesion regression (Fig. 7).

**Figure 7:**
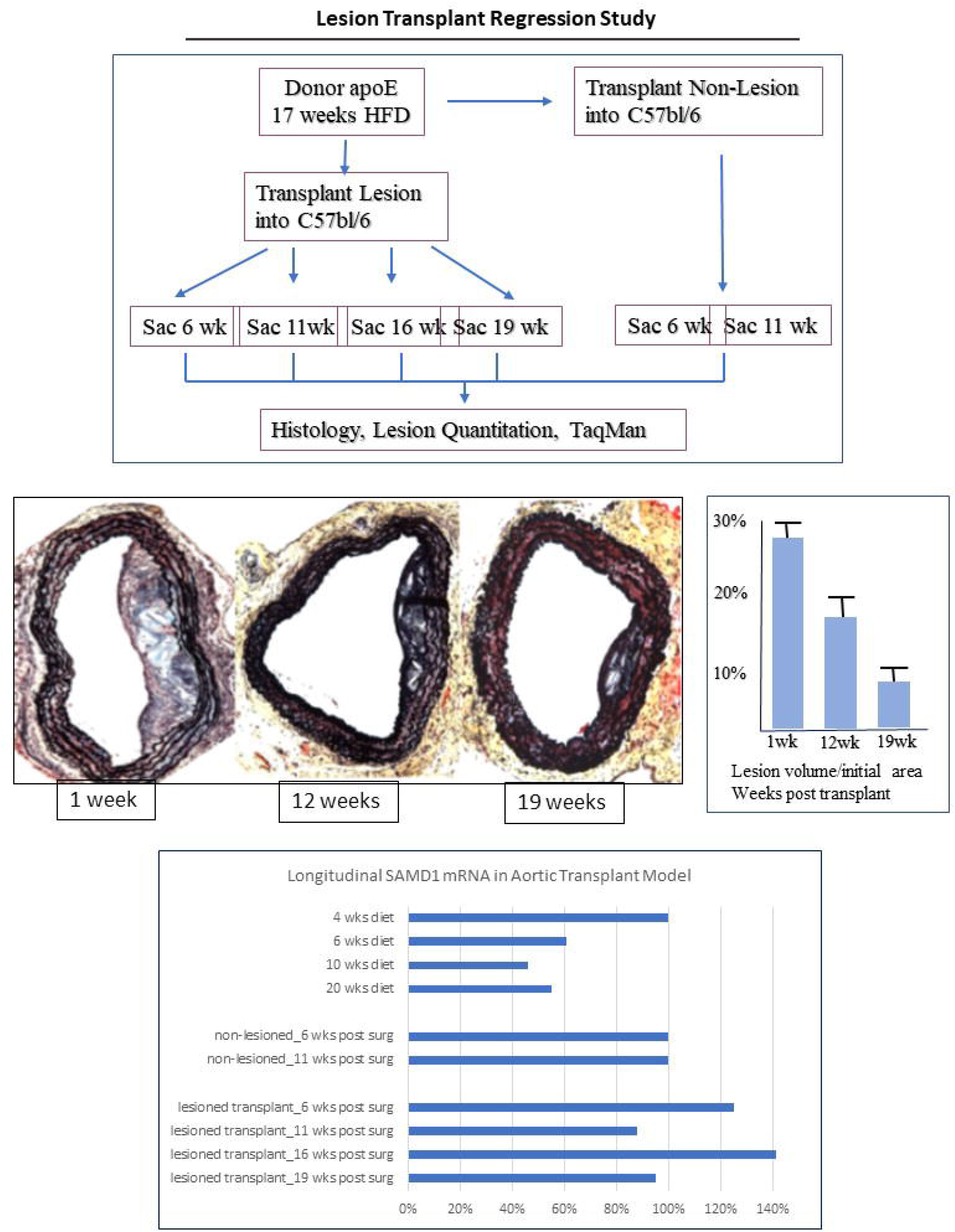
SAMD1 mRNA Expression in Lesion Transplant Regression Model. Aortic sections containing spontaneously formed lesions from apoE-/- mice after 17 weeks on HFD were transplanted into C57BL6 mice on chow diet. Non-lesioned sections were transplanted as controls. Lesion regression was quantified from Movat pentachrome-stained sections, and SAMD1 aortic mRNA levels in the lesions, the transplanted sections, and were measured with Quantigene; results were normalized to non-lesioned sections.

### SAMD1 Non-Lesioned Arterial Localization

Aortas were removed, and stained with GP-abs or monoclonal fabs. ECs from a WT mouse at 10 weeks of age on chow diet stained strongly around the entire circumference, and gauzy faintly stained structures were apparently attached to the luminal surface of many ECs (Fig. 8a); these may be lipid droplets as described by Guyton [15]. All medial SMCs stained in the cytoplasm, and somewhat more intensely at the nuclei. Adventitial staining was seen in scattered cells with various morphologies; in *vasa vasorum*; and in probable *nervi vasorum*. EC staining in an apoE-/- mouse after 3 days on HFD was discontinuous, and some ECs were morphologically different, being domed with thicker and rounded stained nuclei (Fig. 8b). The media in the 3-day apoE-/- mouse had substantially less SAMD1 than the WT, cytoplasmic staining was occasionally foamy, and scattered SMCs were unstained. Staining in a C57BL/6 mouse after 6 days on HFD was indistinguishable from the 3-day apoE-/- (Fig. 8c). No lesions were apparent in an apoE-/-sample after 8 weeks on HFD, but staining patterns had evolved; ECs with moderate nuclear staining were located above small, faint, foamy patterns, and ECs that looked morphologically normal stained less strongly than at 3 days (Fig. 8d). Medial staining at 8 weeks was qualitatively fuzzier than at 3 days, but adventitial staining did not appear different. After 20 weeks, a non-lesioned slice from LDLR-/- mice showed the similar features to the 8-week apoE-/-mice at the lumen, but had infrequent medial stain (Fig. 8e). Morphologically varied adventitial cells stained, as did many adipose cells. Negative controls (NCs) showed no staining.

**Figure 8:**
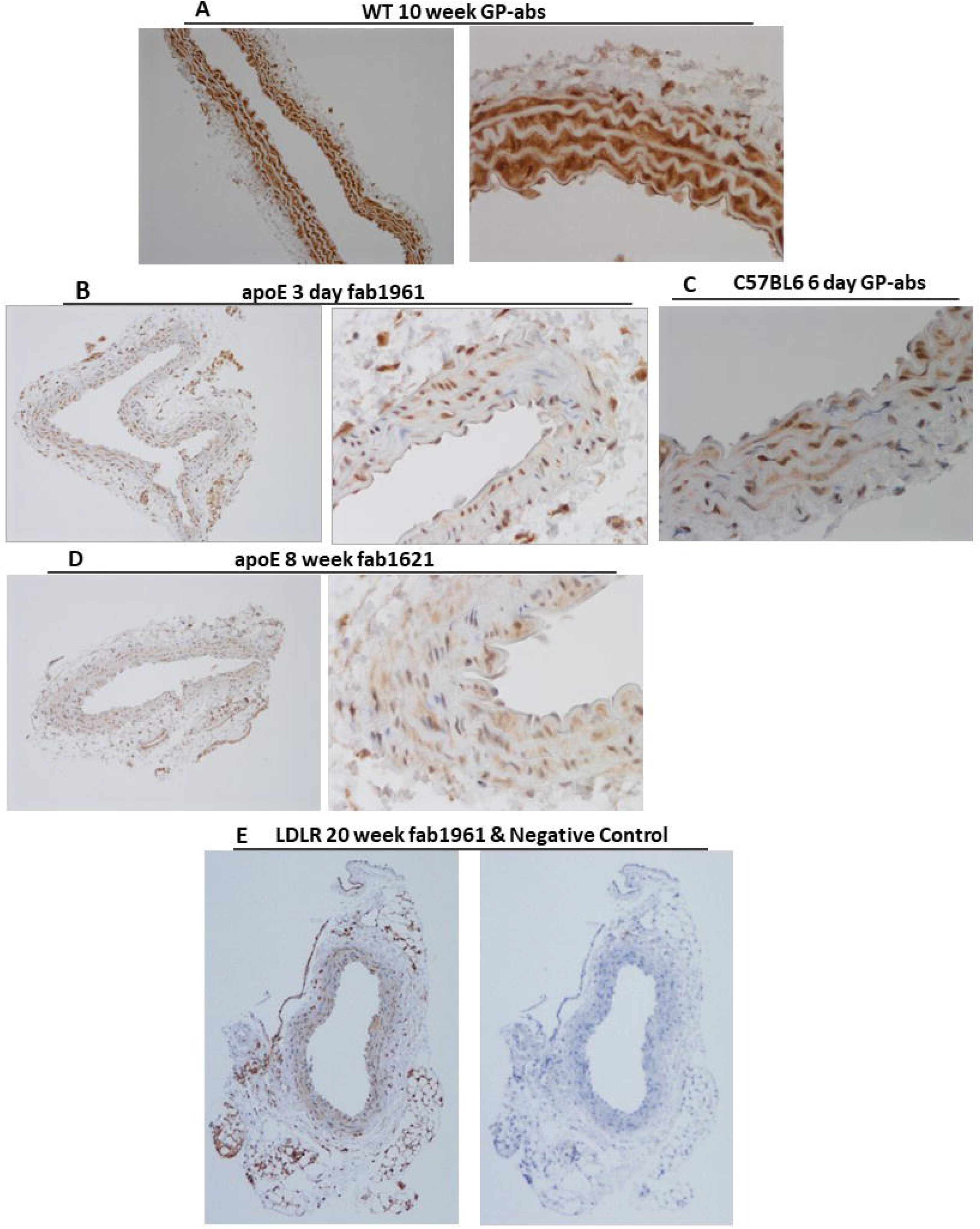
SAMD1 Protein Localization in Non-Lesioned Arterial. Immunohistochemical staining with polyclonal guinea pig anti-rabbit SAMD1 antibodies (GP-abs), and with anti-SAMD1 fab1621 and fab1961. A) Normal aorta from a WT mouse. ECs and medial SMCs stained around the entire circumference; medial SMCs stained somewhat more intensely at their nuclei than cytoplasm. Scattered cells with various morphologies stained in the adventitia. Vasa vasorum were demarcated by SAMD1 stained cells, and SAMD1 stained a probable nervi vasorum. B) Slice from an apoE mouse after 3 days on HFD, with a higher magnification image. SAMD1 is present in all layers of the artery, from endothelial through adipose. C) Slice from a C57BL6 mouse after 6 days on HFD; SAMD1 staining is grossly similar to the 3-day apoE. D) Slice from an apoE mouse after 8 on HFD, with higher magnification image. Compared to the 3-day apoE, medial cells have cytoplasmic stain, and some medial nuclei lack stain. E) Slice from an LDLR mouse after 20 weeks on HFD, with negative control. Adipose stain is pronounced in this sample. All four mouse disease models show substantially less SAMD1 staining than the WT, particularly in the media. EC staining is also reduced.

### SAMD1 Distribution in Lesions

An aorta from a LDLR-/-mouse after 45 weeks on HF diet was co-stained with Alexa488-labeled mAb4B6 (green), polyclonal abs to apoB (blue), and abs to Major Histocompatibility Complex class II (MHCII) (red) (Fig. 9); MHCII is sometimes used to delineate cell surfaces, but is more recently understood as an antigen processing and presenting molecule expressed by multiple cell types, with upregulation indicating immune responses. ECs stained at the cell surface for SAMD1, which was inconsistently colocalized with MHCII and apoB (Fig. 9a). Just inside the cell membrane, SAMD1 stain surrounded structures resembling lipid rafts, but could be caveolae, or small intracellular lipid droplets. Some of these SAMD1-stained structures are immediately adjacent to punctate MHCII. ECs have punctate intracellular MHCII staining, and some ECs have possible nuclear SAMD1 stain. The intima of a lesion from the same aorta consisted of foam cells of various sizes, and smaller cells of unknown types (Fig. 9b). SAMD1 stained the cell surfaces and nuclei of most if not all intimal cells, as well as faintly staining the surfaces of the foamy bubbles that fill foam cells. SAMD1/apoB colocalizations stained strongly in small confluent ovoid foam cells near the lumen; the foamy patterns were increasingly faint as foam cell size increased. MHCII staining is most intense within and around the small foam cells, but also appeared at the cell surface of large foam cells. MHCII was colocalized with SAMD1 at cell surfaces, and in possible endosomes. ApoB was frequently, but not always colocalized with SAMD1 and/or MHCII.

**Figure 9:**
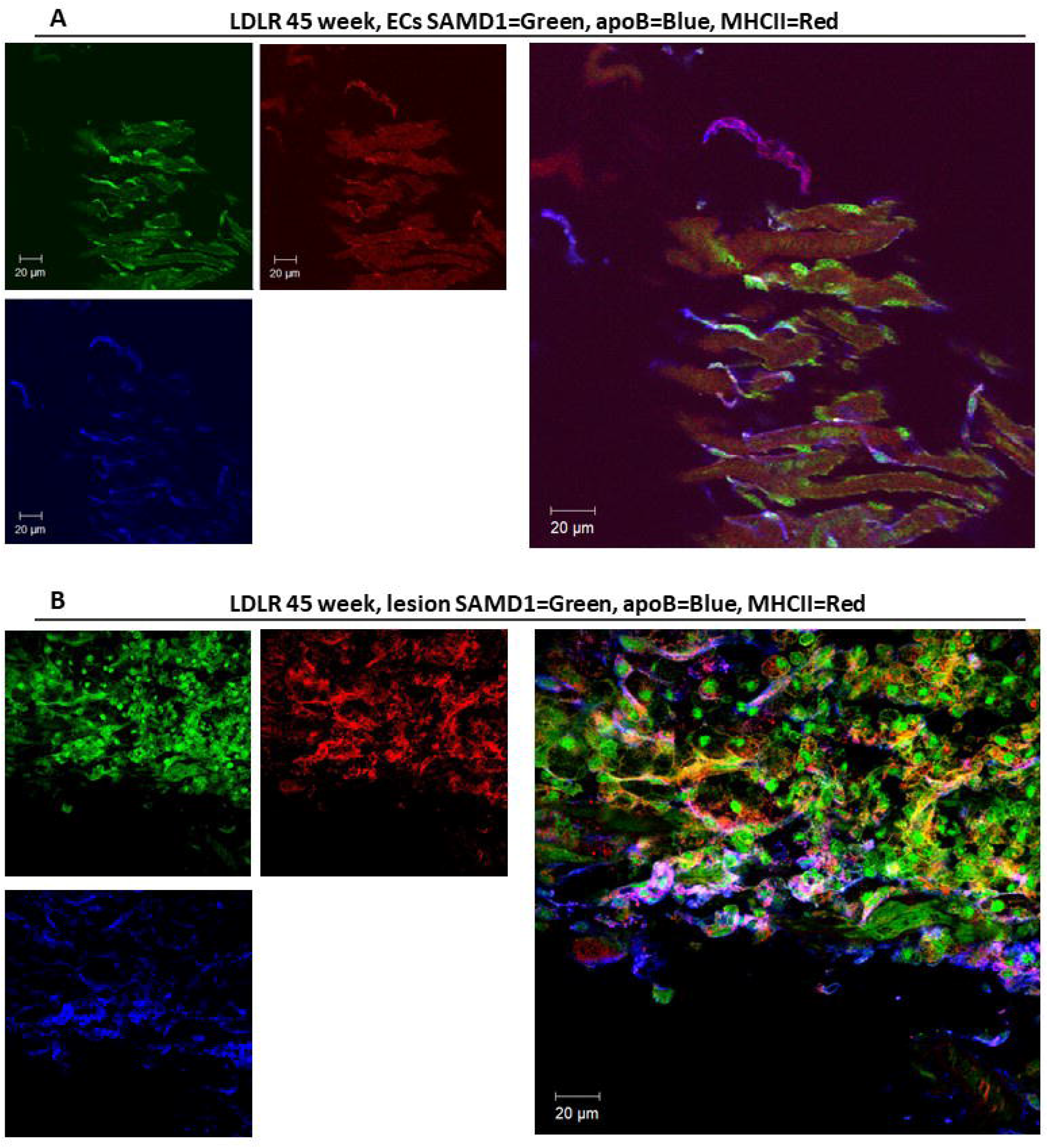
SAMD1 Protein Distribution in Mouse Lesions - Enface Confocal Images. Aorta an LDLR-/- mouse aorta after 45 weeks on HF diet was co-stained with abs to Major Histocompatibility Complex class II (MHCII) (red), polyclonal abs to apoB (blue), and Alexa488-labeled anti-SAMD1 mAb4B6 (green). A) Lumen surface. Just inside the endothelial cell membrane, SAMD1 stain surrounded structures resembling lipid rafts, but could be caveolae, or small intracellular lipid droplets. Some of these SAMD1-stained structures are immediately adjacent to punctate MHCII. ECs have punctate intracellular MHCII staining, and some ECs have possible nuclear SAMD1 stain. MHCII stains cell surfaces in discontinuous stripes, which are inconsistently colocalized with SAMD1 and apoB. Staining for MHCII is faint enough that substantial enhancement of the red signal is required to see punctate intracellular staining, but stronger MHCII staining is seen surrounding the tiny SAMD1-stained foam patches, and at the cell surfaces where it is colocalized with MHCII. B) A small lesion shows SAMD1/apoB/MHCII colocalizations. Most intimal cells are foam cells of various sizes. Staining was particularly intense for SAMD1, apoB, and MHCII in small cells near the lumen, and SAMD1 was colocalized (white-ish) with apoB and MHCII on many but not all cell surfaces. Faint SAMD/MHCII colocalization (yellow/orange) is seen internal to foam cells; and is fainter in the larger foam cells. SAMD1 appeared to be in and immediately around the nuclei of most if not all intimal cells.

Three lesions from a 20-week LDLR-/- suggest lesion development progression (Fig. 10a). A very small lesion consisted of small rounded foam cells and a few spindle-shaped foam cells. SAMD1 stain defined the surfaces of the foamy lipid droplets; most lesional nuclei stained strongly. Beneath the thickest part of the lesion the IEL appeared to have separated. Contralateral to the lesion were several stained lipid-laden macrophage-like cells on the lumen, these cells were adjacent to a thin stained line attached to the IEL, much as reported in bright-field and fluorescent microscopy [16]. Nearby ECs and a dotted line above a foamy structure appeared to be detaching from the IEL. Staining at the lumen in a small 20-week LDLR-/-lesion appeared as a broken dotted and dashed line with stained small rounded nuclei and a few spindle-shaped cells. Mononuclear foam cells beneath the lumen had stained nuclei, but other round nuclei cells did not stain, nor did the few spindle-shaped cells. Unstained lipid accumulations were located against the IEL, and numerous stained adventitial cells were located against the external elastic lamina (EEL). Another 20-week lesion’s intima and adventitia were similar in size and staining patterns, but the lesion protruded because the media had degraded and become hypotrophic.

**Figure 10:**
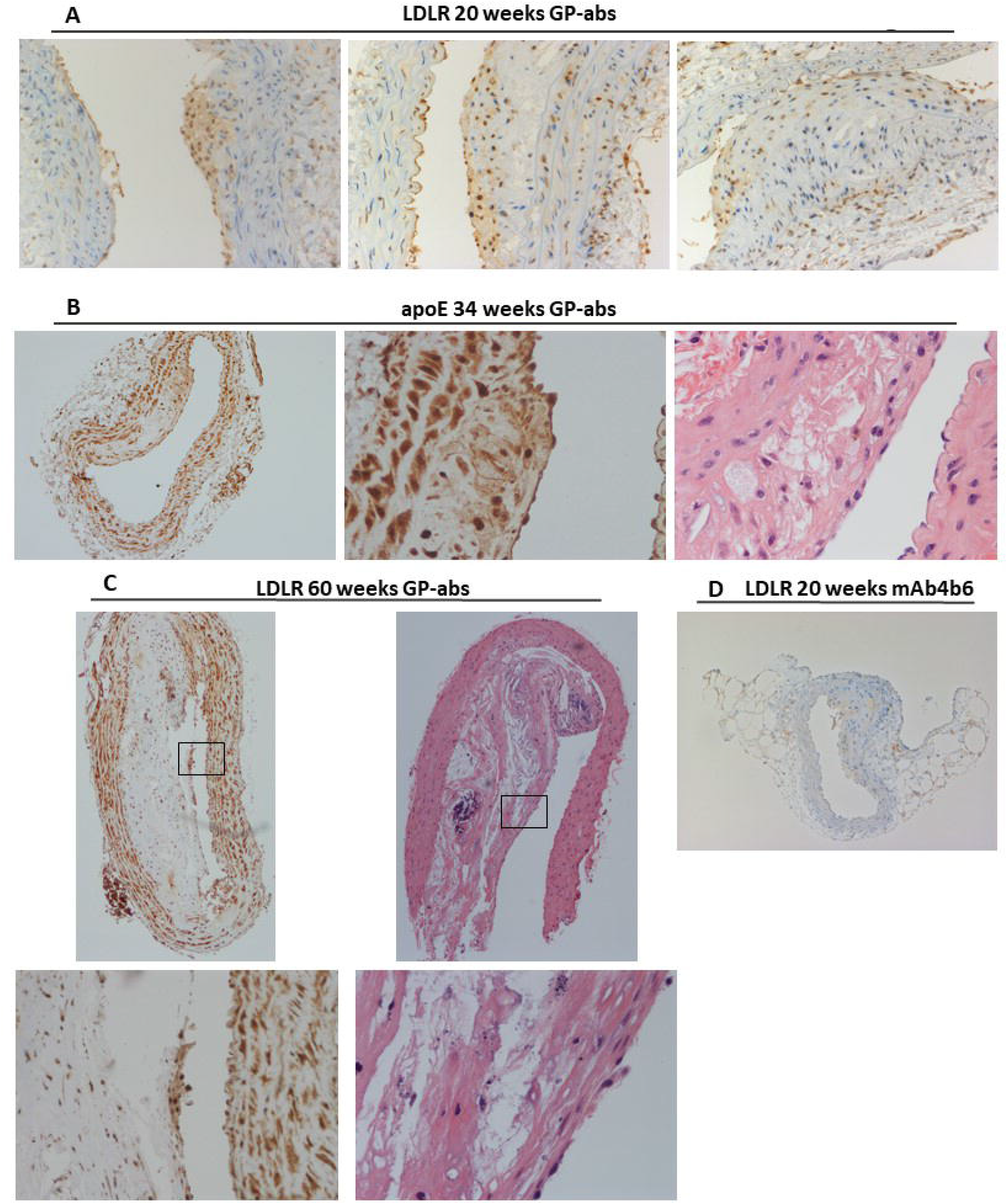
SAMD1 Protein Distribution in Mouse Lesions -IHC Images. Immunohistochemical staining with polyclonal guinea pig anti-rabbit SAMD1 antibodies (GP-abs), and with anti-SAMD1 mAb4B6. a) Three different lesions in LDLR-/- mice after 20 weeks on HF diet. The smallest consisted of small foam cells with stained nuclei, ECs were not morphologically obvious, the IEL was interrupted beneath the thickest part of the lesion, and medial cells had unusually long thin nuclei and may have been dividing. Contralateral to this lesion, an earlier consequence of high LDL is seen at the lumen, with a thin band of SAMD1 and cell-sized foamy structures. A larger lesion had developed an intima with small round and ovoid foam cells; SAMD1 stained these at the nuclei, cell surfaces, and faintly stained the surfaces of the foamy intracellular bubbles. At the lumen, scattered ECs, fibrous-appearing strands, and small nuclei stained. Unstained lipid accumulations were adjacent to the IEL. The media included foam cells and stained round nuclei. Clusters of stained cells aligned along the EEL. A more protrusive lesion had ECs with nuclear stain above stained macrophage foam cells; immediately beneath this layer, SMCs and SMC foam cells had capped the lesion. Beneath the capping layer, round and ovoid foam cells of various sizes stained for SAMD1 on surfaces of the foamy droplets; the nuclei of most intimal cells stained. A substantial part of the protrusion was caused by medial hyperplasia beneath a break in the media, where only scattered nuclei stained for SAMD1, these included foam cells with faint stain. Medial lipid accumulations did not stain for SAMD1. Adventitial staining was increased, stained cells were aligned beneath the EEL; round, lobed, stellate, and elongated cells were seen, and some blood cells in the vasa vasorum appeared to stain. B) apoE-/- mouse after 34 weeks on HFD, with higher magnification SAMD1 image and H&E stain. ECs stained strongly around the entire circumferences and SAMD1-stained spindle-shaped cells capped lesions. Towards the lesion shoulder, the surfaces of lipid droplets in small spindle-shaped and oblong foam cells stained faintly; larger rounded foam cells had diffuse faint intracellular stain, but moderate stain at their cell surfaces. Near the lumen beneath the midcap, small ovoid foam cells stained, but below these cells, staining of the surfaces of foamy intracellular bubbles was rare, and H&E stain confirms that the unstained areas are lipid accumulations. Intima nuclei of uncertain cell types stained. Medial stain was nearly at the level of the WT, with nuclei staining somewhat stronger than the cytoplasm. Adventitial staining was pronounced; round, lobed, stellate, and elongated cells were seen, and a dense cluster of strongly stained cells was in the adventitia on the opposite side from the lesion. Capillary cells in the adventitia also stained. Brown adipose tissue stained strongly. C) A large lesion from an LDLR-/- mouse after 60 weeks on HFD. The lumen above had stained ECs and a few short stretches of aggregated rounded cells with intracellular and nuclear stain. Scattered stained SMCs, and unknown cell types, appeared in the intima. Very small dots stained strongly in some fibrotic areas; based on comparison with H&E slices, these may represent netosis or small calcifications. Necrotic cores had developed, but these had minimal SAMD1 staining, except in clusters of calcified material, and a few small patches of faint uniform stain. Medial stain was strong, and an intensely stained possible ATLO lay against the media. D) 20-week LDLR-/- mouse, stained using inhibitor mAb4B6. This mAb under-stains medial cells compared to fabs and polyclonal abs, but show luminal features similar to the other abs. Adipose staining is seen most clearly in hypertrophic cells.

By 34 weeks on HFD, lesions in an apoE-/- mouse after (Fig. 10b) may have stabilized somewhat compared to the 20-week LDLR-/- lesion. Strongly stained ECs were present, stained SMCs capped the lesions, and intimal nuclei of several leukocyte types stained. A few scattered small ovoid foam cells stained, but less frequently than in the 20-week lesions. At the lesion shoulder there was faint stain on the surfaces of lipid droplets in small spindle-shaped and oblong foam cells; larger rounded foam cells had diffuse faint intracellular stain and moderate stain at their cell surfaces. Fibrotic areas and lipid accumulations were forming, but did not stain. The media beneath the lesion contained breaks, but medial SMCs stained more strongly than at 20 weeks.

60 weeks of HFD in LDLR-/- mice produced large lesions with extensive fibrotic areas, small necrotic cores, lipid pools, and micro-calcifications, as seen by H&E stain (Fig. 10c). A non-sequential slice showed luminal SAMD1 in a few short stretches of round cells, mingled with elongated cells, and stained fuzzy material surrounding clusters of cells. Staining deeper in the intima was limited to scattered cells with nuclear stain, a few small areas of uniform stain in necrotic cores, and intense staining of a cluster of cells that appeared to be calcific. Medial stain was similar to the 34-week samples. The strongly stained cluster of cells adjacent to the media may be an ATLO.

Fragments of brown and white perivascular adipose tissue (PVAT) remained attached to some samples (Fig. 10d). Some PVAT samples stained, others did not; mAb and fab inhibitors, GP-abs, mAbs, and anti-mouse polyclonal fabs all stained sequential slices from PVAT samples that did stain. Staining in a given sample was not uniform - some adipose cells stained, others didn’t; however, the stained cells appeared hypertrophic.

### Lesion Initiation

SAMD1 staining in 20, 34, 45, and 60-week lesions appears to present a continuing progression of LDL retention, foam cell growth, and foam cell necrosis, but SAMD1 localization patterns during lesion initiation (Fig. 11) appears to involve different cell types and/or behaviors than in lesions. A 16-week LDLR-/-mouse appears to have subendothelial stain and foamy medial SMCs in a non-lesioned slice, but much stronger luminal and medial staining in a possible incipient lesion, including foamy patterns at the lumen, and punctate stain in the elastic lamina (Fig. 11a). Three distinct luminal localizations were seen in one slice from an 8-week apoE-/-mouse (Fig. 11b). Clusters of cells with a mix of nuclei shapes enveloped in lacy SAMD1 stain appeared to be attached to the IEL, with no ECs above or below, adjacent to an interruption in the IEL; no cells were above a thin line of strongly stained dots, which are likely platelets [17], that lay directly on the IEL to the left of the cluster, and nearby rounded cells and foamy accumulations were unstained. The lacy SAMD1 enveloping the cell clusters appears to be a mix of aggregation and foamy patterns. Medial stain seemed further decreased, but adventitial stain seemed increased. In Fig. 11b (and Fig. 10a), missing ECs could have been shed into the lumen [18], and platelets may adhere where ECs are lost [19], or ECs were possibly been lost in slide prep. Slice Fig. 11c from the 8-week apoE mouse showed a foamy pattern similar to slice Fig. 11b, but with an intact endothelial layer, and punctate SAMD1 in the media.

**Figure 11:**
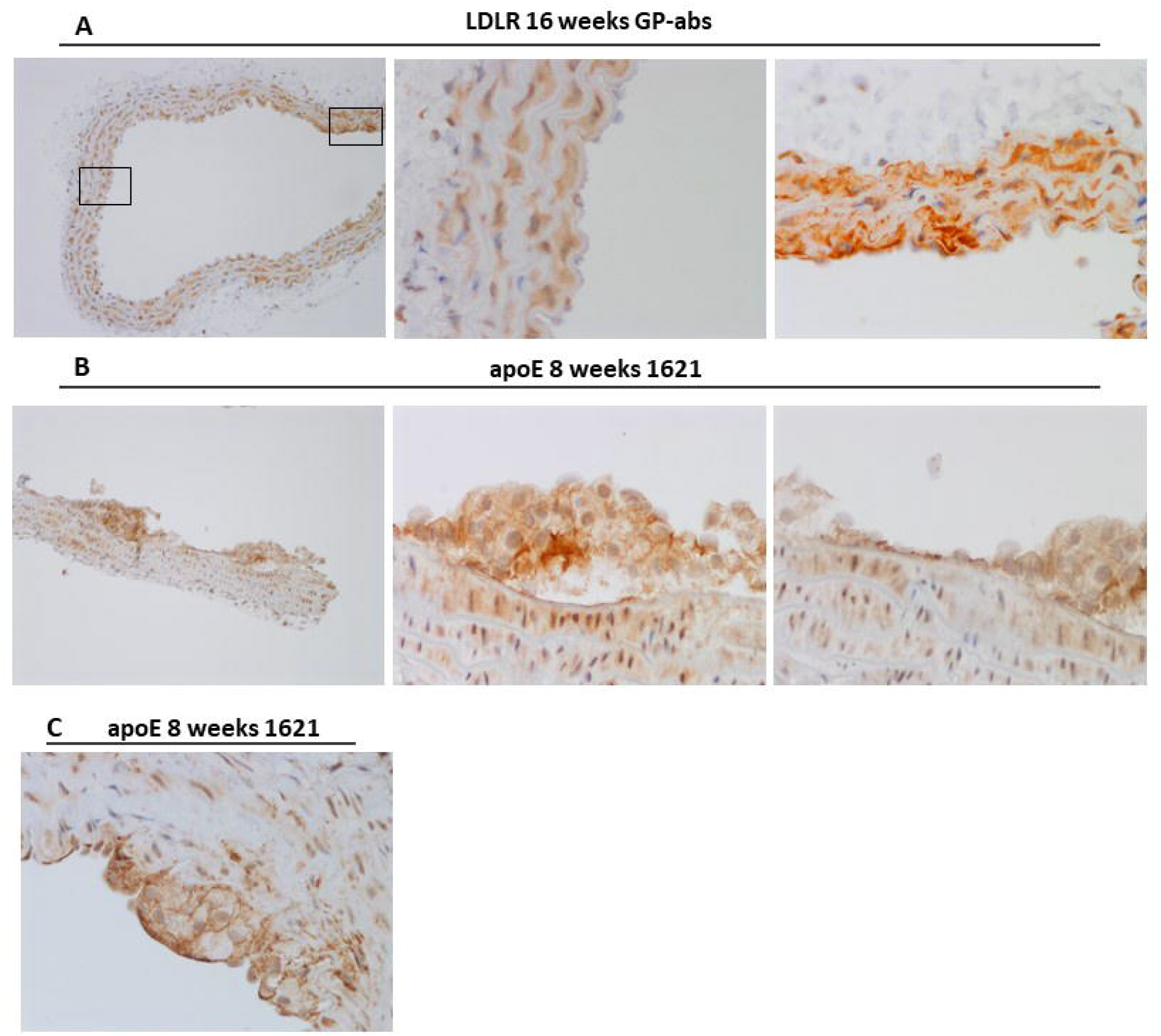
SAMD1 Protein Localization in Lesion Initiation. A) Aorta form an LDLR mouse after 16 weeks on HFD. Subendothelial stain and foamy medial SMCs appear in a non-lesioned area, but much stronger luminal and medial staining in a possible incipient lesion, including foamy patterns at the lumen, and punctate stain in the media. B) Three distinct SAMD1 localizations were seen at the lumen of an 8-week apoE mouse. Dense subendothelial stain apparently prior to the entrance of leucocytes; A cluster of 6-8 subendothelial mononuclear foam cells adjacent to several more individual mononuclear foam cells; punctate medial stain; stained and unstained long thin medial cells. Adjacent to an interruption in the IEL were clusters of cells with a mix of nuclei shapes, enveloped in lacy SAMD1 stain, appeared to be attached to the IEL, with no ECs above or below. No cells were above a thin line of strongly stained dots that lay directly on the IEL to the left of the cluster, and nearby rounded cells and foamy accumulations were unstained. The lacy SAMD1 enveloping the cell clusters appears to be a mix of aggregation and foamy patterns. Medial stain seemed further decreased, but adventitial stain seemed increased. C) A different slice from the 8-week apoE mouse showed a foamy pattern similar B), but with an intact endothelial layer, and punctate SAMD1 in the media.

### Inhibitor Lesion Targeting

Confocal microscopy, MRI, and ultrasound were used to confirm that anti-SAMD1 antibodies would attach to SAMD1 *in vivo*. Cross-section and en face images of a lesioned artery from a 52-week apoE-/- mouse 24 hours after injection with Alexa488-labeled mAb4B6 show patterns similar to IHC, indicating successful antibody penetration of the vessel (Fig. 12a). Inhibitor fab1961, labeled with nano-bubbles, was injected into ApoE-/- mice with carotid injury lesions. Numerous green blebs, ranging from less than .05mm up to .15mm, were seen in the lesion; this is a typical size for lesional lipid droplets (Fig. 12b). Anti-mouse-SAMD mab4B6 was duel-labeled with gadolinium (Gd) micelles and rhodamine, then injected into 56-week -old ApoE-/- mice and 56-week-old LDLR-/- mice (Fig. 12c). Gd is used as a contrast agent in MRI, and rhodamine is used as a red fluorescent tag for confocal microscopy. MRI normalized arterial vessel wall enhancement (%NENH) in the ApoE-/- mice was over 80% at 24 hours; competitive inhibition with unlabeled mAb4B6 reduced enhancement by half. In the LDLR-/- mice, enhancement was over 150%, suggesting some differences between mouse models. *Ex vivo* confocal images of the enhanced areas were taken 24 hours after injection to confirm targeting; rhodamine labeled mAbs (red) and CD68 (green) were colocalized (yellow) in the lesion, showing uptake by CD68-expressing cells.

**Figure 12.**
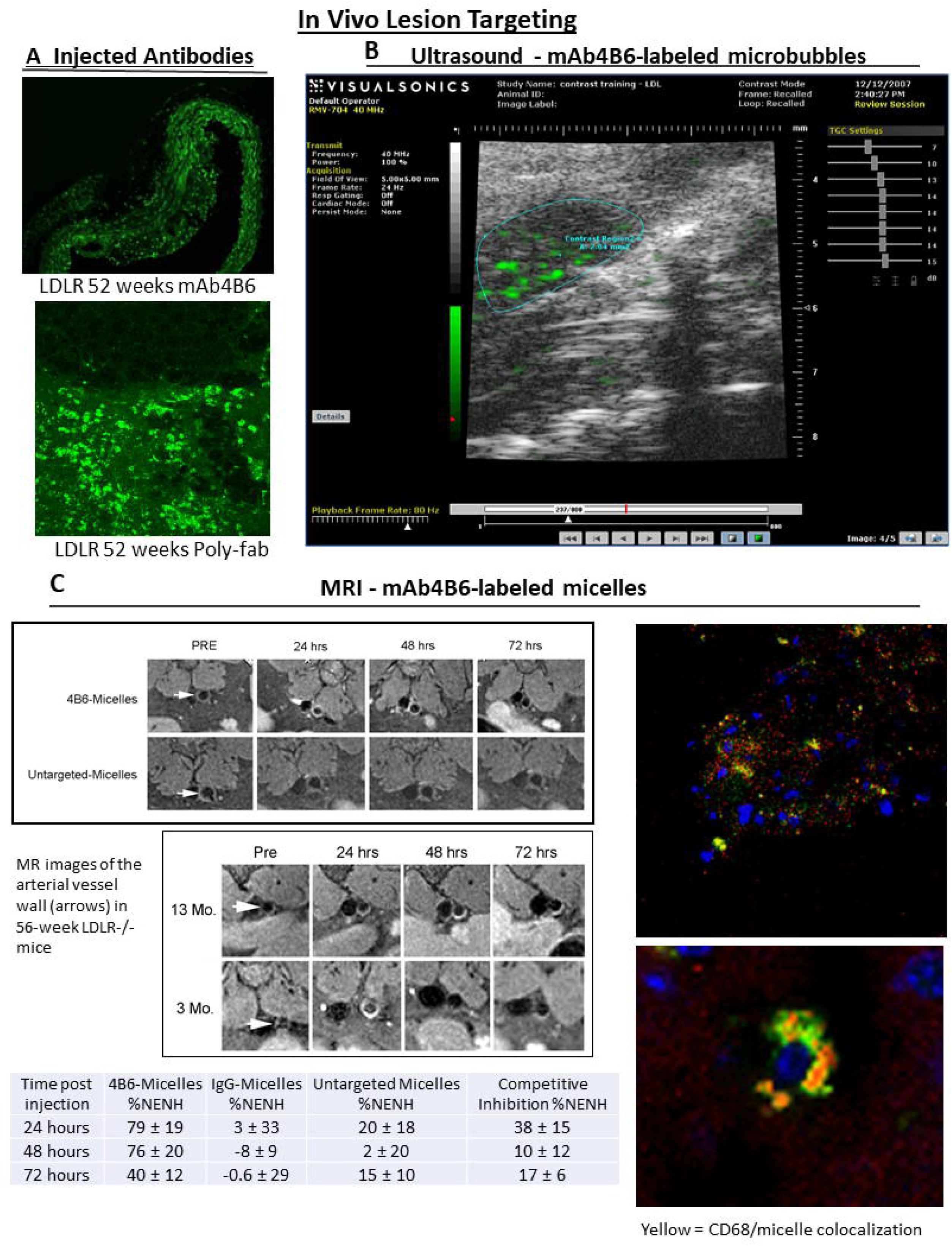
In Vivo Lesion Targeting of Anti-SAMD1 Antibodies. A) Enface and cross section images of an artery from an LDLR-/- mouse 52 weeks on HFD taken 24 hours after in vivo injection of SAMD1/apoB binding inhibitor mAb 4B6 conjugated to Alexa488 (green), showing inhibitor penetrates lesional cells. B) An apoE-/- mouse on HFD had a carotid artery ligated to induce lesions. Two weeks later, microbubbles labeled with fab1961 were injected, and ultrasound imaging performed on a UHF VisualSonics small animal research machine. Numerous green blebs ranging from less than .05mm up to .15mm can be seen in the ligated section. C) mAb 4B6 was covalently linked to the surface of gadolinium micelles, and rhodamine (red) was added as a fluorescent label. Untargeted-micelles (i.e., no antibody present) were used as controls. Representative MR Images of the abdominal arterial wall of 56-week apoE-/- mice following administration of 4B6-micelles and untargeted micelles (0.075 mmol Gd/Kg), as a function of time post injection. Arrow indicates the aorta. - Representative MR images of the arterial vessel wall (arrow) in LDLR-/- mice as a function of mouse age. - In vivo competitive inhibition studies were performed to test the specificity of the 4B6-micelles for LDL epitopes in the arterial wall. Age matched apoE-/- mice (n=3) were intravenously administered 0.5 mg/mouse of free 4B6 immediately prior to the administration of 4B6-micelles (0.075 mmol Gd/kg and 0.26 mg 4B6) and MR enhancement was compared to MR enhancement obtained after the administration of 4B6-micelles (n=3). Percent-normalized enhancement (%NENH) using 4B6 labeled micelles, relative to muscle, was determined for the aortic vessel wall and reflect the percent change in the contrast-to-noise ratios (CNR) obtained pre and post injection. Competition of free 4B6 prior to intravenous injection of 4B6-micelles resulted in ∼2-fold reduction in %NENH. - Confocal images of the arterial vessel wall of an apoE-/- mouse that had been in vivo injected with rhodamine labeled 4B6-micelles (red) and ex vivo stained with abs to CD68 (green). Yellow shows colocalization of CD68 with 4B6-micelles; cell nuclei are blue.

### SAMD1 Inhibitors and Lesion Growth

Spontaneous lesion and carotid injury models were used to test if blocking SAMD1 could impact lesion size. LDLR-/-mice and apoE-/- mice were put on HFD for 12 weeks. The right carotids were ligated one week prior to treatment with two doses of 10mg/kg peg-Fab1621 over a three-day period. Mice were then injected with 125I-TC-labelled human LDL and sacrificed after 24 hours. Right-injured and left-uninjured carotid arteries were removed as well as the thoracic aorta. The carotids and 2mm sections of lesioned and non-lesioned areas from the thoracic aorta were analyzed using autoradiographic techniques. Carotid injury lesions from untreated apoE-/- mice retained twice as much I125TC-labeled human LDL injury lesions from treated apoE-/- mice. The effect size was about half as large in treated LDLR-/- mouse carotid injury lesions, while spontaneous left carotid and thoracic lesions from these mice showed a less than 10% treatment effect (Fig. 13a).

**Figure 13:**
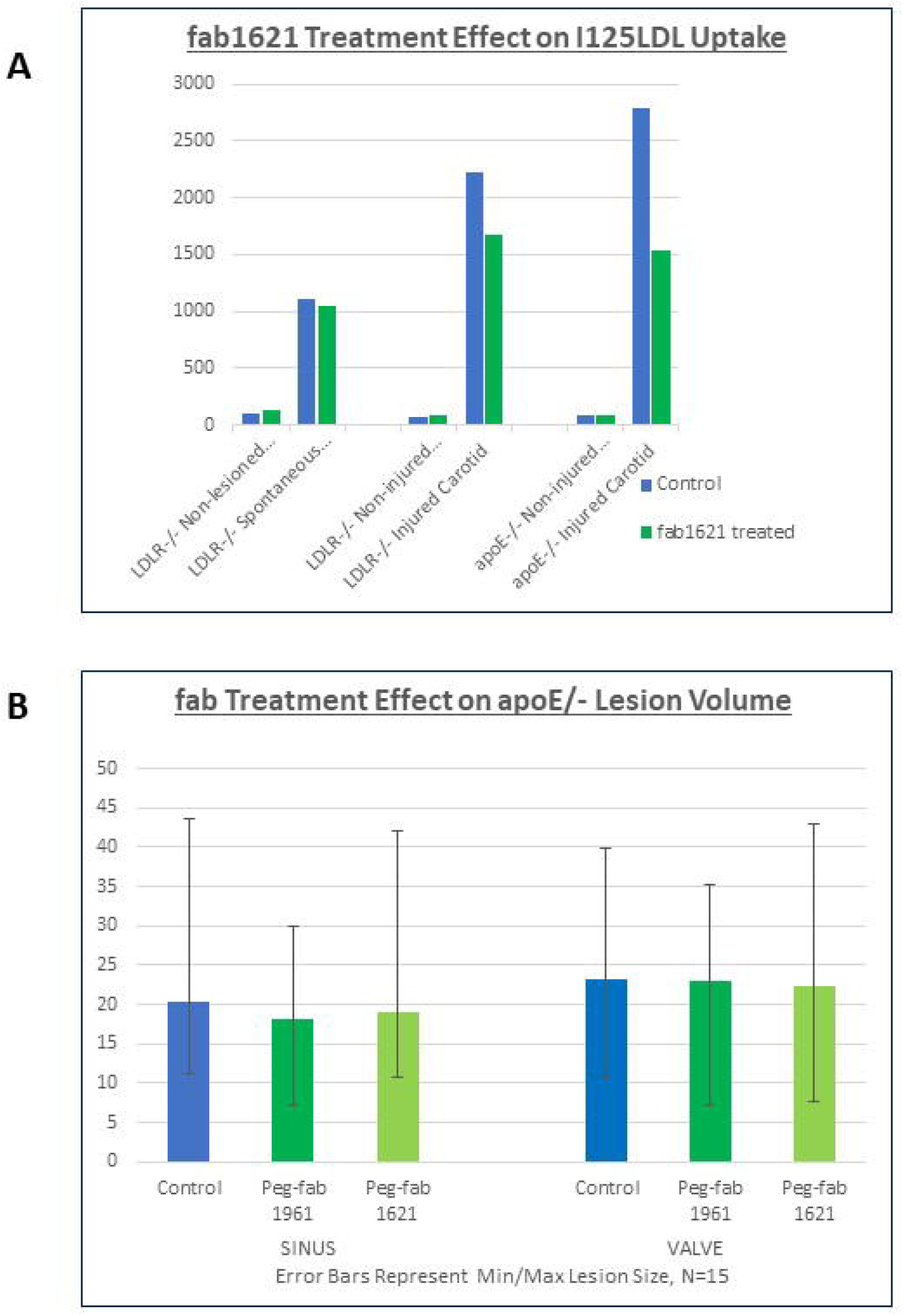
Effects of SAMD1/LDL Binding Inhibitors on Lesion Initiation. Spontaneous lesion and carotid injury models were used to test whether SAMD1/LDL binding inhibition could affect lesion size. A) Carotid injury lesions took up substantially less 125I-TC-labelled human LDL after treatment with two doses of 10mg/kg peg-Fab1621 over a three-day period. LDLR-/-mice and apoE-/- mice were put on a modified western diet for 12 weeks to induce spontaneous aortic lesions. The right carotids were injured one week prior to inhibitor treatment and sacrificed 24 hours after the second dose. Right-injured and left-uninjured carotid arteries were removed as well as the thoracic aorta. The carotids and 2mm sections of lesioned and non-lesioned areas from the thoracic region were analyzed using autoradiographic techniques. Right carotid lesions generated by ligation in treated apoE-/- mice with retained 50% less I125TC-labeled human LDL, and 27% less in lesions in treated LDLR-/- mice. Spontaneous left carotid and thoracic lesions showed less than a 10% difference in LDL uptake. B) 10% treatment effect was seen on spontaneous valve and sinus lesion area from apoE-/- mice after 12 weeks on modified western diet {1.25% chol} when treated with 10mg/kg peg-Fab1621 or peg-Fab1961every other day. Smaller effects were seen with peg-Fab1961. The results did not achieve statistical significance (N=15 for each treatment group and controls). Consistent with that result, I125TC had minimal uptake difference in the apoE spontaneous lesions in these A) same mice; data not shown. Lesion size varied by over a factor of four within each group; the smallest and largest lesions from each group are shown as error bars.

In contrast to the carotid injury lesions, but consistent with LDL uptake in spontaneous lesions, only 10% treatment effect was seen on spontaneous valve and sinus lesion area from apoE-/- mice after 12 weeks on modified western diet {1.25% chol} when treated with 10mg/kg peg-Fab1621 every other day. The results did not achieve statistical significance (N=15 for each treatment group and controls). Effects were smaller with peg-Fab1961. Lesion size varied by over a factor of four within each group; the smallest and largest lesions from each group are shown as error bars (Fig. 13b).

## Discussion

This paper presents a continuation of our research into atherogenic retention of LDL, beginning with finding intense focal LDL accumulation at the edges of regenerating endothelial islands in ballooned rabbit arteries [20]; irreversible focal LDL accumulation for at least 40 h [21]; accumulation of a synthetic peptide comprising apoB residues 1000–1016 showing a focal pattern indistinguishable from that of whole LDL [22]; focal accumulation of LDL in human atherosclerotic plaques suggesting that the specificity of LDL accumulation that we saw in rabbits also occurred in human disease [23]; and co-localization of LDL and SAMD1 in human lesions supporting a specific mechanism for focal LDL binding in atherosclerotic lesions [2]. In this study, we provide data to support the hypothesis that SAMD1 not only plays a role intimal retention of LDL in mouse models of atherosclerosis, but SAMD1 staining patterns change throughout the arterial vessel as lesions develop.

### SAMD1/LDL Binding

Inhibition studies with full-length and truncated versions of SAMD1 suggest a possible SAMD1/LDL binding domain are to the N-terminal, possibly around a.a. 344, or to a different folding of some cross domain. Since some mAbs to the mouse fragment inhibit, the inhibitors could occupy that differential domain (about a.a. 193-237), or the folding is different between the CMT mouse fragment and the Biosite fragment. The binding model is probably complicated, since ELISAs showed somewhat weaker binding to the human fragment that starts at a.a.344. However, two mAbs that were weak inhibitors attached tightly to a fragment from the 3’end of human SAMD1 that started at a.a. 344, so cross-domain SAMD1-LDL binding is possible. Although the binding location on apoB is uncertain, SAMD1 might bind to LDL by changing its conformation and returning to the initial conformation after binding is complete, similar to HIV protease. This is supported by the observation that SAMD1/apoB binding inhibitors effectively attached to both solo SAMD1 and SAMD1/apoB complex. The fact that PEGylation improved the IC50 of the fabs suggests that steric interference is important to inhibition. It is not clear how the inhibitors function *in vivo*, given that the *in vitro* results suggest the inhibitors don’t dissociate an established SAMD1-LDL complex, instead, it implies ongoing expression of SAMD1 in growing lesions such that inhibitor-accessible SAMD1 epitopes are available prior to LDL binding.

In intimal lesions, SAMD1 appears to bind LDL irreversibly. It seems likely that SAMD1/apoB complex seen in the ECM is the source for SAMD1/apoB complex on the surface of the foamy lipid droplets that fill foam cells. This is in accordance with reports that the foamy pattern is caused by progressive loss of apo B from LDL leaving aggregated cholesteryl droplets [24], [25], which means that recognizable epitopes of SAMD1 and apoB have not dissociated in spite of partial catabolism. The presence of SAMD1/apoB in foam cells means that the complex has been ingested by the cell, which presumably only happens if it is recognized as antigenic.

### Treatment Effect

Lack of treatment effect in early lesions challenges our other observations that strongly support SAMD1’s role in lesional LDL retention and foam cell formation. Our earlier attempts to slow lesion initiation with peptide inhibitors of SAMD1/LDL binding, and with immunization against SAMD1 [26], did not produce statistically significant results (unpublished observations). Studies have shown that inhibiting the LDL/proteoglycan interaction slows early lesion growth, but has no impact on lesion size or volume by 26-35 weeks because another retention mechanism becomes dominant as lesions progress [27]. The current observations from mice at 20, 34, and 45 weeks suggest that SAMD1 is involved in lesion growth via apoB-Lps retention and ingestion, and thus inhibitors might have shown treatment effect after 30-40 weeks. Substantial treatment effect in the carotid injury model may be due to the combination of stimuli that leads to very rapid SMC-rich lesion development. Only 2 days after injury, there is a 60% reduction in the number of medial SMCs [28]; medial SMCs rapidly dedifferentiate, migrate to the intima, and proliferate such that at two weeks, about 80% of cells in the developing neointima are SMC, many of these have differentiated to macrophage-like foam cells [29], [30]. We thus hypothesize that SAMD1 expressed by intimal SMCs is the LDL binding molecule [2], [3], that gradually comes into effect in spontaneous lesions as intimal SMCs migrate and proliferate, starting between weeks 8 and 12 [31].

The extreme variations in lesion sizes seen in cloned mice suggests unknown factors are important in atherogenesis. These variations may be related to the lack of consistent results seen in our exploration of SAMD1 mRNA expression in cell culture. SMC phenotype modulation appears to be an important part of lesion development, and, in addition to phenotype modulation in cultured cells being driven by various chemical factors, culturing conditions and media, and cell density, plated SMCs are not phenotypically stable [14]; in the case of SMCs, which are phenotypically plastic, small differences in otherwise identical cell culture runs may produce large differences in outcomes.

### SAMD1 as a Probe/Marker

Early atherosclerosis research was primarily focused on intimal lesions; it is now increasingly recognized that the entire arterial vessel participates [32]. Immunostaining for SAMD1 may be useful for connecting changes that occur across the artery, and our data suggests SAMD1 may have roles in atherosclerosis beyond LDL retention. Variations in SAMD1 stain appear to correlate with phenotype changes in multiple cell types, and it is possible that differences seen in stained nuclei reflect SAMD1’s epigenetic functions. Cell surface staining of foam cells and ECs may be from antigen presentation. Staining for SAMD1 and SAMD1/apoB colocalizations appears to allow tracking the fate of apoB-lipoproteins from the lumen to apoptotic foam cells to calcifications. Note that multiple intimal cells have phenotypes that bind and process apoB-Lps, and it is difficult to distinguish between SMCs, macrophages, dendritic cells, neutrophils, or other cells that may be athero-related. SAMD1 stain also suggests involvement with platelets and netosis in lesions.

### SAMD1 in the Endothelial Layer

SAMD1 in the endothelial layer appears to be involved in lipid uptake and processing. By definition, LDL must transit the EC layer for subendothelial retention to occur. ECs have at least four mechanisms for transporting LDL [33]; SAMD1 appears to be involved in one or more EC transport mechanisms, as well as EC antigen presentation. The colocalized SAMD1/apoB seen at parts of EC cell surfaces (Fig. 9a) suggests SAMD1/apoB complex is being formed at endocytosis. The lipid droplets or caveolae in ECs surrounded by SAMD1 stain, often with apoB immediately adjacent, sometimes with MHCII immediately adjacent, suggests endosomal/lysosomal processing of SAMD1/apoB complex. Further, ECs above subendothelial lipid accumulation can become lipid-filled [34], and oxLDL has been reported in EC foam cells [35]; SAMD1 staining on the surfaces of foamy lipid droplets in ECs suggests that SAMD1 is involved in their creation. ECs are antigen presenting cells [36] and leukocyte responses to EC antigen presentation are antigen-specific. MHCII/apoB complexes are reported on antigen-presenting cells in mouse lesions [37], possibly expressed in response to oxLDL [38]; our observation of MHCII/apoB colocalizations at certain parts of EC cell surfaces implies that some apoB-containing molecules, without SAMD1, are being presented as antigens that will attract leukocytes. Other parts of EC cell surfaces have SAMD1/MHCII colocalizations without apoB, implying antigen presentation of a SAMD1-related molecule. These distribution patterns suggest that SAMD1 is involved in EC LDL uptake and or transport, as well as antigen presentation.

This raises questions about LDL antigenicity. SAMD1/apoB complex is, and/or becomes, sufficiently modified to be recognized as antigenic; SAMD1/apoB and SAMD1/CD68 colocalizations in probable lysosomes/endosomes in human and mouse lesions, and SAMD1/apoB on the surfaces of foam cell lipid droplets means that SAMD1/apoB complex has been ingested and processed. In mouse models, LDL antigenicity may be induced by luminal interaction with ECs, during EC LDL transport, and/or during subendothelial retention [34]. A recent study reports that LDL binding to SAMD1 on SMC surfaces induced LDL oxidation [3], so the SAMD1/apoB complex may be antigenic soon after formation. Stimulated ECs recruit and retain platelets, and release extracellular vesicles [39]. The function of cytoplasmic SAMD1 in our cultured ECs is unknown, but the circular colocalizations of SAMD1/MHCI adjacent to nuclei may be lysosomes (Fig. 4). SAMD1 staining (Figs 10a, & 11) is seen on what may be platelets and extracellular vesicles which carry RNA, proteins, and lipids as a form of extracellular communication [40]; the source and function of SAMD1 here is unknown, but may be involved in antigen presentations and/or communications which signal and attract leukocytes [41]. Since lesion formation is eccentric, it is interesting that in (Fig. 10a) this staining, which is contralateral to a small lesion, includes several apparent monocytes.

### SAMD1 in the Intima

In the lesioned mouse intima, SAMD1’s appearance is consistent with a role in LDL retention as per the extended R2R theory. Our confocal microscopy shows: SAMD1 is highly expressed by most nuclei of cells in the lesion; in cell culture, SAMD1 can be translocated to the ECM of SMCs; SAMD1 and apoB are colocalized it the ECM of most cells in the lesion; SAMD1 and apoB are colocalized on the surfaces of foamy lipid droplets; SAMD1 and SAMD1/apoB stain decreases in intensity as foam cell size increases, to the point that some intracellular foamy patterns can be seen with H&E do not stain for SAMD1.

SAMD1 upregulation and translocation to the ECM are probably caused by stress, injury, and disease signals from other cells that can trigger rapid SMC phenotype modulation [42]. SAMD1 mRNA expression was inconsistently increased in cultured SMCs when stressed, and results suggested that signaling between ECs and SMCs is relevant. It seems likely that stressors cause differentiated SMCs to secrete SAMD1 to the ECM with actin, as seen in our confocal images of cultured SMCs (Fig. 4). SAMD1, once in the ECM and at the cell surface, is accessible to bind perfusing LDL. VSMC phenotype modulation can not only be induced by SAMD1 up- and down-regulation, but increasing and suppressing SAMD1 expression on the outside surface of the VSMC cell membrane increases and decreases retained LDL, oxLDL, and foam cell formation [3].

Intimal staining for SAMD1 hints that the fate of apoB-lipoproteins can be tracked from lumen to lipid pools. Although the timing and means of LDL’s becoming antigenic in the intima are not yet agreed on, no other mechanism for unregulated LDL uptake is well supported. That implies that SAMD1/apoB complex ingested from the ECM is the source of SAMD1/apoB complex in foam cells. The decreasingly intense SAMD1 and SAMD1/apoB stain seen in foam cells of increasing size appears to continue to the point that some intracellular foamy patterns that can be seen with H&E do not stain for SAMD1. These distribution patterns suggest that the larger foam cells develop from ongoing retention of apoB-Lps by SAMD1, followed by ingestion and catabolism of SAMD1/apoB complex. CD68+ cells such as foam cells hydrolyze the cholesterol esters (CE) in LDL to free cholesterol because only unesterified (free cholesterol, (FC)) can be effluxed from the cells to extracellular cholesterol acceptors [43], [44]. It seems that catabolism continues until the SAMD1 and apoB are extensively digested as a part of the process during which the CE has been hydrolyzed to FC. FC lipid accumulations imply competent foam cell processing of ingested SAMD1/apoB-Lps, and thus explain the absence of SAMD1 in mouse lipid pools. This is consistent with the free-cholesterol-rich vesicles that form mouse model lipid accumulations after foam cell necrosis [44], [45], [46].

In lesions of 60-week mice, SAMD1 appears to stain two types of calcifications commonly seen in advanced mouse lesions: osteogenic cells and calcifying matrix vesicles (Fig. 10). SMCs can change phenotype to become osteoblast-like and chondrocyte-like; SMCs and macrophages can release matrix vesicles that are colocalized with cholesterol [47]. The small dots seen in fibrotic areas that stain blue in the H&E and also stain strongly for SAMD1, are probably calcific matrix vesicles, some of which may be aggregating [48], [49]. It would seem that these, like lipid pools, are another advanced fate for SAMD1/LDL complex.

SAMD1 staining at the cell surface of morphologically different cells in the intima may denote phenomena different from extracellular LDL retention, such as ingestion of SAMD1 via endocytosis/phagocytosis/transcytosis, or presentation of a SAMD1 epitope as an antigen. Antigen presentation by multiple cell types, including macrophages, SMCs, dendritic cells, and B lymphocytes found in atherosclerotic lesions is widely reported. Our confocal images of SAMD1, apoB, and MHCII colocalizations at cell surfaces in the intima are consistent with reports of apoB/MHCII complexes on antigen-presenting cells in mouse lesions, possibly expressed in response to oxLDL [36], [38]. Antigen-presenting cells continuously display self-peptides from apoB/LDL [50]; SAMD1 is likely a part of this complex. Additional research is needed to understand if SAMD1’s appearance at the cell surface is an artifact of its irreversible complexing with apoB, or if it is involved in the creation of the presented antigen, and/or if SAMD1 binding makes LDL antigenic. Foamy SAMD1 localization patterns in the earliest subendothelial lipid accumulations, prior to SMC arrival, suggests that SAMD1 could be involved in leukocyte uptake, and/or processing, of antigenic apoB-lipoproteins. SAMD1/apoB colocalization could show retention in the basal lamina, but this cell surface staining might also occur early in ingestion, or could be due to antigen presentation; it is not possible to tell with the resolutions we used.

Monocytes enter the intima and convert to macrophages of different types; medial SMCs migrate to cap lesions, proliferate, and modulate to macrophage-like phenotypes and osteogenic phenotypes; dendritic cells activate and ingest lipids; T-cells convert to atherosclerosis specific proinflammatory types as disease progresses [13]. The nuclei of most intimal cells stain for SAMD1, but the cell types are not identifiable by our methods, so it is not clear if SAMD1 expression here is related to cell phenotype changes.

### SAMD1 in the Media

Reduced SAMD1 staining was seen in medial SMCs from young mice born with dyslipidemia/hypercholesterolemia, as compared to the WT (Fig. 8). We presume that the SAMD1 expression in WT medial SMCs is normal, and the disease models are born in a state of responding to abnormal lipids. This is consistent with reported downregulation of medial SMC genes, adaptive immune responses, and lipid accumulations that are apparent 10 days after conditional knockout of the apoE-/-gene in mice [51]. It is also consistent with a recent finding that both up- and down-regulation of multiple pathways was seen in SAMD1 KO ES cells, including up-regulation of muscle cell migration [5]. In contrast, another study found that upregulation rather than downregulation of SAMD1 caused phenotype changes in VSMCs [3]. In apoE-/-mice after 16 weeks on HFD, medial SMC markers are reduced and some medial SMCs are expressing CD68 [52], suggesting additional phenotype changes that correlate with reduced SAMD1 staining. As the mice aged and lesions grew, medial SMC staining increased; one possible explanation is that medial SMCs revert to a more stable phenotype as lesions stabilize.

### SAMD1 in the Adventitia

Changes in SAMD1 staining in the adventitia may be related to proliferation of immune cells and/or cell phenotype changes. The adventitia senses and responds to inflammatory, hormonal, and environmental stresses and controls arterial inflammation response, resolution, lesion formation, remodeling, and creation of neointima [53], [54]. The majority of adventitial cells are fibroblasts; other cell types include: resident progenitor cells, dendritic cells, macrophages, T-cells, B cells, mast cells, vasa vasorum and lymphatic endothelial cells and nerve cells [54]. In WT and young mice, scattered adventitial cells of various morphologies have nuclear and/or intracellular SAMD1 stain. Qualitatively, the number of stained cells was increased, and aligned along the EEL near lesions in the 20 week mice; this may be related to the proliferation of adventitial fibroblasts that drives neointima formation [55]. There were no obvious differences in adventitial staining between the 20-week and 45-week mice. By 60 weeks, the adventitia appears to have degraded or atrophied, so staining data was not useful. Adventitial Tertiary Lymphoid Organ (ATLOs) can be formed during nonresolving peripheral inflammation, typically adjacent to lesions [56], and are probably formed by transdifferentiated SMCs [57], [58]. Although intense SAMD1 staining in a probable ATLO suggests T-cell recognition of SAMD1 in atherosclerosis, it is possible that SAMD1’s appearance in antigen presentation is an artifact of SAMD1’s irreversible binding to apoB.

### SAMD1 in Perivascular Adipose Tissue

Limited adipose SAMD1 staining data was available, and was insufficient to find correlation with disease state. HFD in mice provokes a strong immune response in PVAT, resulting in infiltration of immune cells [59]. Based on samples with valid negative controls (NCs), PVAT staining for SAMD1 was in patterns similar to reports for CD68 [60], MAC1 [61], and NOTCH2 [61], noting that both brown and white PVAT stained. Although no adipose staining was seen in NCs that used irrelevant antibodies, the small set of adipose samples had to be reduced because brown and white PVAT was stained in NCs for four out of 16 samples that used guinea pig serum or mouse anti-gst supernatant for the secondary only controls. This suggests that in certain disease states Fc receptors in PVAT are expressed that are either non-specific (mouse serum control) and/or cross-species (guinea pig serum control).

## Conclusion

In conclusion, this is the first study characterizing SAMD1 expression, localizations, and colocalizations in mice. SAMD1 appears to be important in atherosclerotic LDL retention and antigen presentation; an approximate timepoint for SMC translocation of SAMD1 becoming the dominant mechanism of LDL accumulation is 18-20 weeks. Foam cells appear to competently degrade SAMD1/apoB complex before they become apoptotic/necrotic [62] and release lipids and cell debris which build up deeper in the thickening lesion. Nuclear SAMD1 may imply epigenetic regulatory activities; additional research is required to clarify how these are connected to the changes in nuclear and cytoplasmic SAMD1 staining seen in endothelial, medial, adventitial, and adipose layers as lesions develop. Our data suggests that SAMD1 may be a useful probe for improving understanding of lesion development.

## Methods and Materials

### Immunohistochemistry

We generated guinea pig polyclonal antibodies to a rabbit SAMD1 fragment expressed by e coli that started at a.a. 238 and continuing to the 3’ end (GPSBPO2), as previously described [2]. Full-length human SAMD1 was produced by Cell & Molecular Technologies (CMT, Phillipsburg, NJ, now Invitrogen, Carlsbad, CA) in HEK293 cells using proprietary methods. The human SAMD1 coding sequence (1617bp) was redesigned such that the codon usage was optimized without changing the amino acid sequence. Full-length human SAMD1 was expressed at high levels in transfected HEK293 cells. However, attempts to extract the protein by cell lysis in TBS/Triton X-100 resulted in incomplete recovery of soluble protein. Extraction of the protein by sequential homogenization in low salt and high salt buffers using a Teflon-glass homogenizer recovered about 10% of the protein; storage in D-PBS w/o Ca/Mg at dry ice temperatures prevented precipitation and aggregation. Partial mouse SAMD1 cDNA was expressed as a ∼45 kDa protein in the cytoplasm of transfected HEK293 cells. In addition to the 45 kDa protein, a ∼30 kDa protein was also expressed. This fragment also carries the FLAG tag consistent with the notion that it is a C-terminal degradation product of the 45 kDa protein. Anti-FLAG Western blot suggests that the two forms are expressed in roughly equimolar amounts. Both forms were soluble in TBS/Triton X-100 extracts. Biosite (Biosite/Alere/Abbott, San Diego, CA) transfected E coli to produce SAMD1, but E coli were only able to express a human C-terminus fragment that started at aa344 and went to the end at aa538.

A&G Pharmaceutical, Inc. (Columbia, MD) generated mouse and human mAbs as described [63], and performed SAMD1/LDL binding affinity assays. 96 well ELISA plates were coated with 200ng per well of LDL freshly extracted LDL provided by Pfizer (Groton, CT) (note that commercially available LDL from four different vendors was non-functional as tested by three of our partners). Our earlier work [2] was reproduced using SAMD1 rabbit 3’ from E.coli, added at a range from 0-1ug. 1:1000 of anti-mouse SAMD1 serum was used for primary Ab, 1:2000 of HRP-goat anti-mouse IgG (H+L) Ab was used for secondary Ab, TMB was the substrate. These methods were next used to establish LDL binding to full-length human SAMD1 and the above mouse SAMD1 fragment.

Biosite (Biosite/Alere/Abbott, San Diego, CA) made fully-human fab-fragments to full-length human SAMD1, and fab1961 by phage display; monoclonal fabs were selected by ELISA for affinity to full-length human SAMD1. SAMD1/LDL binding behavior was explored using ELISA competitive inhibition. The first ELISA used two blocks of fabs in a sequential format wherein SAMD1 was allowed to bind to freshly extracted LDL provided by Pfizer (Groton, CT) on the plate first, before washing and adding fab, resulting in a maximum signal that was dependent on antibody concentration and affinity for SAMD1; i.e., no inhibition was observed. The second ELISA was performed by mixing fab and SAMD1 before adding the mixture to LDL on the plate. The two assays were compared, with the lower signal being interpreted as inhibited binding. The final concentration of SAMD1 was the same in both sequences, 500 ng/ml, allowing comparison between the assays. Note that age of LDL seemed to slightly affect total signal and inhibition; older LDL seemed to generate a higher signal but lower inhibition.

<u>**Sequential** SAMD1 & Antibody Steps</u>

a. LDL Ag: 5ug/ml in NaP buffer
b. Block with CD8
c. SAMD1: 100ng/ml, 0
d. Antibody: 0.5ug/ml
e. GAM K-AP: 1:300

<u>**Pre-mixed** SAMD1 & Antibody Steps</u>

a. LDL Ag: 5ug/ml in NaP buffer
b. Block with CD8
c. SAMD1: 100ng/ml, 0
d. Antibody: SAMD1 Ag (100ng/ml,0) + Antibody (0.5ug/ml) - pre-mixed 1hr.
e. GAM K-AP: 1:300

Pfizer (Groton, CT) used a Biacore Surface Plasmon Resonance machine to determine SAMD1/LDL binding kinetics; LDL as “ligand” was bound to the sensor and SAMD1 as “analyte” was channeled over the surface. The ligand and analyte were then reversed, with SAMD1 bound to the sensor and LDL channeled over the surface.

Immunohistochemical studies using anti-SAMD1 antibodies, MHCI, MHCII, DAPI, H&E, and alpha smooth muscle actin and immunofluorescence studies were performed by the Brigham & Women’s Hospital Pathology Lab (Boston, MA). Note that IHC and IF staining for SAMD1 was reliable with formalin fixed-paraffin embedded, but not using frozen fixation; however, in vivo injection of both anti-SAMD1 fabs and mAbs labelled with Alexa488 provided adequate signals in dissected-out tissue.

An ultrasound imaging study was performed by Pfizer (Groton, CT): apoE-/- carotid arteries were ligated to induce lesions; microbubbles were labeled with mouse anti-human-SAMD1 monoclonal fab1961; ultrasound imaging was performed on a UHF VisualSonics small animal research machine.

MRI studies and immunohistochemical staining procedures using anti-SAMD1 antibodies and CD68, were performed by the Mount Sinai School of Medicine (New York, NY): A total of twenty-three 12-month-old male apoE^-/-^ mice were used: n=8 for untargeted-micelles, n=8 for mAb4B6-micelles, and n=4 IgG-micelles, and n=3 for competitive inhibition studies. In addition, six 6 LDLR-/- mice were also scanned: n=3 at 12 months of age and n=3 at 5 months of age). Nine 12-month-old male wild type (WT) mice were also used: n=3 for untargeted-micelles, n=3 for 4B6-micelles and n=3 for IgG-micelles. All strains of mice were of a C57BL/6 background. ApoE^-/-^ and LDLR-/- mice were placed on a high-cholesterol diet (0.2% total cholesterol, Harlan Teklad, Madison, Wisconsin) ad libitum beginning at 6 weeks of age. WT mice were maintained on a normal murine diet (Research Diets, Inc., New Brunswick, NJ). The ethics committee at Mount Sinai approved all of the animal experiments. The mechanism of 4B6-micelle uptake was evaluated in ex vivo murine J774A.1 macrophages (8^th^ generation). All macrophages were plated into 12-well plates in Dulbecco’s modified Eagle’s medium (DMEM) containing fetal calf serum. All macrophages were incubated with MDA-LDL (5 µg/ml) for 24 hours at 37° C. The wells were then washed in fresh DMEM and the macrophages were exposed to untargeted-micelles (n=2), mAb4B6-micelles (n=2), or IgG-micelles (n=2) (all at 1 mM Gd). Saline controls (n=2) were also included. In a second 12 well plate, the MDA-LDL fed cells were exposed to the micelles after an initial incubation period of the micelles with LDL (1 hour at 37° C). Micelles were pre-incubated with LDL (Calbiochem, 5 µg/ml) in order to determine if the mAb4B6-micelles enter the cell only after binding with LDL. All cells were then incubated for an additional 24 hours at 37° C.

### RNA and cDNA Measurements

RNA and cDNA measurements were performed by the Brigham & Women’s Hospital Pathology Lab (Boston, MA), and by Pfizer (Groton, CT). Synthesis of cDNA (ThermoScript RT-PCR System (Invitrogen Cat 11146-016)). RT-PCR was performed in a MyiQ single-color real-time PCR system (Bio-Rad Laboratories, Inc., Hercules, CA) at the Brigham, and QuantiGene (Thermofisher) at Pfizer. Total ribonucleic acid (RNA) from 250,000 cells was reverse transcribed by Superscript II (Invitrogen) according to the manufacturer’s instructions. Quantitative PCR was performed with SYBR green PCR mix (primers for SAMD from QIAGEN N.V.) and analysis performed with StepOne Software (ThermoFisher, Applied Biosystems). mRNA levels were normalized to glyceraldehyde 3-phosphate dehydrogenase (GAPDH) mRNA levels. SAMD1 sample and controls cDNA were synthesized (ABI high-capacity cDNA Archive Catalog No. 4387406). Total 2ugs RNA/100ul cDNA synthesis reaction and 1ul cDNA per Taqman Reaction. Mouse SamD1 Assay – Mm01261650_g1. Mouse GAPDH endogenous control-(Applied Biosystems™ 4352339E. Catalog No. 43-523-39E).

### Cell Culture Studies

Cell culture SAMD1 expression studies were performed by Pfizer (Groton, MA),

1. Preparation - cells and medium:
  Human Aortic Endothelial Cells (HAoE C): Clonetics, cc-2535
  Human Aortic Smooth Muscle Cells (HAoSMC): Clonetics, cc-2571
  EGM-2 BulletKit for HAoEC: Clonetics, cc-3162
  SmGM-2 Bulletkit for HAoSMC Clonetics, cc-3182
  24 mm Transwell Insert: 0.4 um Pore polycarbonate membrane Transwell Insert (3412)
2. Culture of Cells
  Seeding density for all 3 type cells is 10000 or 20000 cells/cm2, and time points for culture are 24, 48 and 72 hrs.
    1 × 105 or 2×105 cells in 2 ml medium are added to a well (9.6 cm2) of 6 well plates;
    5×104 or 1×105 cells in 1.5 ml medium are added to an insert (4.67cm2).
  Co-culture of EC/SMC mixture:
    5×104 EC in 1 ml of EC medium
    5×104 SMC in 1 ml of EC medium, mix them.
  Co-culture of EC/SMC with insert:
    1 × 105 or 2×105 SMC in 2 ml of EC medium
    5×104 or 1×105 EC in 1.5 ml of EC medium
  Experiment Groups:
    HAoSMC alone:
    HAoEC alone:
    HAoSMC/HAoEC mixture:
    HAoEC insert/SMC bottom:
3. RNA expression levels were quantitated using the probes and the Panomics QuantiGene kit from:
  HAoSMC:
  HAoEC:
  HAoSMC/HAoEC mixture:
  HAoSMC/HAoEC mixture + mechanical injury by pipette tip scratch
  SMC bottom
  EC insert
  SMC bottom + EC injury
  EC insert +injury:

Cell culture studies as above, with the addition of chemical stressors, and two different SAMD1 probes, were performed by Brigham & Women’s Hospital Pathology Lab (Boston, MA). Note that the study was complicated by the fact that most cell types express SAMD1, making it difficult to design an effective RT-PCR negative control. Therefore, the lab explored expression of SAMD1 protein in cultured cells using immunofluorescent labelling in confocal microscopy.

- Mouse aortic smooth muscle cells were plated at low confluence. Cells were co-stained for SAMD1 using Alexa488-labeled mAb4B6, MHCI (red), DAPI (blue), alpha smooth muscle actin.
- Mouse heart endothelial cells were plated at low confluence. Cells were co-stained for SAMD1 using Alexa488-labeled mAb4B6, MHCI (red, DAPI (blue).

### Animal Studies

Mouse lesion regression transplant study was performed by Pfizer (Groton, MA). Aortic sections containing spontaneously formed lesions from apoE-/- mice after 17 weeks on HFD were transplanted into C57BL6 mice on chow diet. Non-lesioned sections were transplanted as controls. Lesion regression was quantified from Movat pentachrome-stained sections. SAMD1 aortic mRNA levels in the lesions, the transplanted sections, and non-lesioned sections were measured with Quantigene; results were normalized to non-lesioned sections.

Mouse lesion SAMD1 uptake studies were approved and performed by Pfizer (Groton, MA). LDLR-/-mice and apoE-/- mice were put on a modified western diet for 12 weeks to induce spontaneous aortic lesions. Five days pre-procedure, mice were injected with 125I-TC-labelled human LDL as described [64]. The right carotids were injured one week prior to inhibitor treatment and sacrificed 24 hours after the second dose; treatment consisted of two doses of 10mg/kg peg-Fab1621 over a three-day period. Right-injured and left-uninjured carotid arteries were removed, as well as the thoracic aorta. The carotids and 2mm sections of lesioned and non-lesioned areas from the thoracic region were analyzed using autoradiographic techniques.

Mouse lesion growth treatment studies were approved and performed by the Brigham & Women’s Hospital Pathology Lab (Boston, MA). apoE-/- mice were placed on modified western diet (1.25% cholesterol), and treated with 10mg/kg peg-Fab1621 or peg-Fab1961 every other day for 12 weeks. Atherosclerotic lesions at the aortic valve and sinus were analyzed as described [65]. Briefly, the aorta was dissected out and fixed. Serial 10 um thick cryosections of valve and sinus were collected for a distance of 400 um. These sections were stained with Oil Red O and hematoxylin. The lipid staining areas on 25 sections each for valve and sinus were determined by light microscopy and summed as a proxy for lesion volumes.

## Acknowledgements

We thank Dr. Andrew Lichtman (Brigham & Women’s Hospital, Boston, MA) for guiding discussions, critical inputs, unwavering intellectual support, and managing members of his lab throughout this research, including Dr. Margaret Tarrio, who performed mouse treatment studies, light and confocal microscopy, and various assays, and Dr. Israel Gotsman, for confocal and light microscopy. We thank Dr. Karen Saeboe (Mount Sinai School of Medicine, now with Invicro) for MRI and related confocal studies; Dr. Joe Corvera (A&G Pharmaceutical, Inc., Columbia, MD) who developed and tested mAb inhibitors of SAMD1/LDL binding; Dr. Roland Tacke (CMT, Phillipsburg, NJ) for expression and purification of SAMD1; Dr. Louis Kerr (Woods Hole Oceanographic Institute) for his expertise with the confocal microscopy; Dr. Gunars Valkirs (now Biosite /Alere/Abbott, San Diego, CA) for guiding the development and testing of fab inhibitors of SAMD1/LDL binding; and Dr. Walter Gilbert (Boston, MA) for laying out and explaining all the steps needed to develop and test SAMD1/LDL binding inhibitors.

## Bibliography

[1] K. J. Williams and I. Tabas, “The response-to-retention hypothesis of early atherogenesis,” Arterioscler. Thromb. Vasc. Biol., vol. 15, no. 5, pp. 551–561, May 1995, doi: 10.1161/01.atv.15.5.551.

[2] A. M. Lees, A. E. Deconinck, B. D. Campbell, and R. S. Lees, “Atherin: a newly identified, lesion-specific, LDL-binding protein in human atherosclerosis,” Atherosclerosis, vol. 182, no. 2, pp. 219–230, Oct. 2005, doi: 10.1016/j.atherosclerosis.2005.01.041.

[3] S. Tian et al., “The miR-378c-Samd1 circuit promotes phenotypic modulation of vascular smooth muscle cells and foam cells formation in atherosclerosis lesions,” Sci Rep, vol. 11, no. 1, Art. no. 1, May 2021, doi: 10.1038/s41598-021-89981-z.

[4] B. Stielow, C. Simon, and R. Liefke, “Making fundamental scientific discoveries by combining information from literature, databases, and computational tools – An example,” Computational and Structural Biotechnology Journal, vol. 19, pp. 3027–3033, Jan. 2021, doi: 10.1016/j.csbj.2021.04.052.

[5] B. Stielow et al., “The SAM domain-containing protein 1 (SAMD1) acts as a repressive chromatin regulator at unmethylated CpG islands,” Science Advances, vol. 7, no. 20, p. eabf2229, May 2021, doi: 10.1126/sciadv.abf2229.

[6] E. Engelen et al., “Proteins that bind regulatory regions identified by histone modification chromatin immunoprecipitations and mass spectrometry,” Nat Commun, vol. 6, p. 7155, May 2015, doi: 10.1038/ncomms8155.

[7] A. D. Rouillard et al., “The harmonizome: a collection of processed datasets gathered to serve and mine knowledge about genes and proteins,” Database (Oxford), vol. 2016, 2016, doi: 10.1093/database/baw100.

[8] A. Lachmann et al., “Massive mining of publicly available RNA-seq data from human and mouse,” Nat Commun, vol. 9, no. 1, p. 1366, 10 2018, doi: 10.1038/s41467-018-03751-6.

[9] R. Villaseñor et al., “ChromID identifies the protein interactome at chromatin marks,” Nat Biotechnol, pp. 1–9, Mar. 2020, doi: 10.1038/s41587-020-0434-2.

[10] T. C. Huff et al., “Oscillatory cAMP signaling rapidly alters H3K4 methylation,” Life Science Alliance, vol. 3, no. 1, Jan. 2020, doi: 10.26508/lsa.201900529.

[11] L. I. James et al., “Discovery of a chemical probe for the L3MBTL3 methyllysine reader domain,” Nat Chem Biol, vol. 9, no. 3, pp. 184–191, Mar. 2013, doi: 10.1038/nchembio.1157.

[12] W. Sun et al., “A potential regulatory network underlying distinct fate commitment of myogenic and adipogenic cells in skeletal muscle,” Sci Rep, vol. 7, p. 44133, 09 2017, doi: 10.1038/srep44133.

[13] S. Allahverdian, C. Chaabane, K. Boukais, G. A. Francis, and M.-L. Bochaton-Piallat, “Smooth muscle cell fate and plasticity in atherosclerosis,” Cardiovasc. Res., vol. 114, no. 4, pp. 540–550, 15 2018, doi: 10.1093/cvr/cvy022.

[14] L. M. Shackelton, D. M. Mann, and A. J. T. Millis, “Identification of a 38-kDa Heparin-binding Glycoprotein (gp38k) in Differentiating Vascular Smooth Muscle Cells as a Member of a Group of Proteins Associated with Tissue Remodeling *,” Journal of Biological Chemistry, vol. 270, no. 22, pp. 13076–13083, Jun. 1995, doi: 10.1074/jbc.270.22.13076.

[15] J. R. Guyton and K. F. Klemp, “Early extracellular and cellular lipid deposits in aorta of cholesterol-fed rabbits,” Am J Pathol, vol. 141, no. 4, pp. 925–936, Oct. 1992.

[16] R. Mora, F. Lupu, and N. Simionescu, “Prelesional events in atherogenesis: Colocalization of apolipoprotein B, unesterified cholesterol and extracellular phospholipid liposomes in the aorta of hyperlipidemic rabbit,” Atherosclerosis, vol. 67, no. 2, pp. 143–154, Oct. 1987, doi: 10.1016/0021-9150(87)90274-7.

[17] G. Ed Rainger et al., “The role of platelets in the recruitment of leukocytes during vascular disease,” Platelets, vol. 26, no. 6, pp. 507–520, 2015, doi: 10.3109/09537104.2015.1064881.

[18] D. A. Chistiakov, Y. V. Bobryshev, and A. N. Orekhov, “Neutrophil’s weapons in atherosclerosis,” Experimental and Molecular Pathology, vol. 99, no. 3, pp. 663–671, Dec. 2015, doi: 10.1016/j.yexmp.2015.11.011.

[19] A. Faggiotto, R. Ross, and L. Harker, “Studies of hypercholesterolemia in the nonhuman primate. I. Changes that lead to fatty streak formation.,” Arteriosclerosis: An Official Journal of the American Heart Association, Inc., vol. 4, no. 4, pp. 323–340, Jul. 1984, doi: 10.1161/01.ATV.4.4.323.

[20] A. B. Roberts et al., “Selective accumulation of low density lipoproteins in damaged arterial wall,” J Lipid Res, vol. 24, no. 9, pp. 1160–1167, Sep. 1983.

[21] M. Y. Chang, A. M. Lees, and R. S. Lees, “Time course of 125I-labeled LDL accumulation in the healing, balloon-deendothelialized rabbit aorta,” Arterioscler Thromb, vol. 12, no. 9, pp. 1088–1098, Sep. 1992, doi: 10.1161/01.atv.12.9.1088.

[22] I. L. Shih, R. S. Lees, M. Y. Chang, and A. M. Lees, “Focal accumulation of an apolipoprotein B-based synthetic oligopeptide in the healing rabbit arterial wall.,” PNAS, vol. 87, no. 4, pp. 1436–1440, Feb. 1990, doi: 10.1073/pnas.87.4.1436.

[23] A. M. Lees et al., “Imaging human atherosclerosis with 99mTc-labeled low density lipoproteins.,” Arteriosclerosis: An Official Journal of the American Heart Association, Inc., vol. 8, no. 5, pp. 461–470, Sep. 1988, doi: 10.1161/01.ATV.8.5.461.

[24] H. S. Kruth and B. Shekhonin, “Evidence for loss of apo B from LDL in human atherosclerotic lesions: extracellular cholesteryl ester lipid particles lacking apo B,” Atherosclerosis, vol. 105, no. 2, pp. 227–234, Feb. 1994, doi: 10.1016/0021-9150(94)90053-1.

[25] R. A. van Dijk et al., “Differential expression of oxidation-specific epitopes and apolipoprotein(a) in progressing and ruptured human coronary and carotid atherosclerotic lesions,” J. Lipid Res., vol. 53, no. 12, pp. 2773–2790, Dec. 2012, doi: 10.1194/jlr.P030890.

[26] A. Lees, R. Lees, S. Law, and A. Arjona, “Novel low density lipoprotein binding proteins and their use in diagnosing and treating atherosclerosis,” US20020129388A1, Sep. 12, 2002 Accessed: Sep. 09, 2021. [Online]. Available: https://patents.google.com/patent/US20020129388A1/en

[27] I. Tabas, K. J. Williams, and J. Borén, “Subendothelial lipoprotein retention as the initiating process in atherosclerosis: update and therapeutic implications,” Circulation, vol. 116, no. 16, pp. 1832–1844, Oct. 2007, doi: 10.1161/CIRCULATIONAHA.106.676890.

[28] Kumar Anjali and Lindner Volkhard, “Remodeling With Neointima Formation in the Mouse Carotid Artery After Cessation of Blood Flow,” Arteriosclerosis, Thrombosis, and Vascular Biology, vol. 17, no. 10, pp. 2238–2244, Oct. 1997, doi: 10.1161/01.ATV.17.10.2238.

[29] M. Liu and D. Gomez, “Smooth Muscle Cell Phenotypic Diversity,” Arterioscler. Thromb. Vasc. Biol., vol. 39, no. 9, pp. 1715–1723, Sep. 2019, doi: 10.1161/ATVBAHA.119.312131.

[30] B. P. Herring, A. M. Hoggatt, C. Burlak, and S. Offermanns, “Previously differentiated medial vascular smooth muscle cells contribute to neointima formation following vascular injury,” Vascular Cell, vol. 6, p. 21, 2014, doi: 10.1186/2045-824X-6-21.

[31] A. Misra et al., “Integrin beta3 regulates clonality and fate of smooth muscle-derived atherosclerotic plaque cells,” Nat Commun, vol. 9, no. 1, Art. no. 1, May 2018, doi: 10.1038/s41467-018-04447-7.

[32] H. W. Kim, H. Shi, M. A. Winkler, R. Lee, and N. L. Weintraub, “Perivascular Adipose Tissue and Vascular Perturbation/Atherosclerosis,” Arteriosclerosis, Thrombosis, and Vascular Biology, vol. 40, no. 11, pp. 2569–2576, Nov. 2020, doi: 10.1161/ATVBAHA.120.312470.

[33] X. Zhang, W. C. Sessa, and C. Fernández-Hernando, “Endothelial Transcytosis of Lipoproteins in Atherosclerosis,” Front. Cardiovasc. Med., vol. 0, 2018, doi: 10.3389/fcvm.2018.00130.

[34] M. Simionescu, “Implications of Early Structural-Functional Changes in the Endothelium for Vascular Disease,” Arteriosclerosis, Thrombosis, and Vascular Biology, vol. 27, no. 2, pp. 266–274, Feb. 2007, doi: 10.1161/01.ATV.0000253884.13901.e4.

[35] A. V. Sima, C. S. Stancu, and M. Simionescu, “Vascular endothelium in atherosclerosis,” Cell Tissue Res, vol. 335, no. 1, p. 191, Sep. 2008, doi: 10.1007/s00441-008-0678-5.

[36] P. Libby, A. H. Lichtman, and G. K. Hansson, “Immune Effector Mechanisms Implicated in Atherosclerosis: From Mice to Humans,” Immunity, vol. 38, no. 6, pp. 1092–1104, Jun. 2013, doi: 10.1016/j.immuni.2013.06.009.

[37] J. Mai, A. Virtue, J. Shen, H. Wang, and X.-F. Yang, “An evolving new paradigm: endothelial cells – conditional innate immune cells,” Journal of Hematology & Oncology, vol. 6, no. 1, p. 61, Aug. 2013, doi: 10.1186/1756-8722-6-61.

[38] S.-H. Choi et al., “SYK regulates macrophage MHC-II expression via activation of autophagy in response to oxidized LDL,” Autophagy, vol. 11, no. 5, pp. 785–795, May 2015, doi: 10.1080/15548627.2015.1037061.

[39] E. Hergenreider et al., “Atheroprotective communication between endothelial cells and smooth muscle cells through miRNAs,” Nat Cell Biol, vol. 14, no. 3, pp. 249–256, Mar. 2012, doi: 10.1038/ncb2441.

[40] M. Gherghiceanu et al., Part One: Extracellular Vesicles as Valuable Players in Diabetic Cardiovascular Diseases. IntechOpen, 2019. doi: 10.5772/intechopen.85225.

[41] P. Han et al., “Platelet P-selectin initiates cross-presentation and dendritic cell differentiation in blood monocytes,” Science Advances, vol. 6, no. 11, p. oeaaz1580, Mar. 2020, doi: 10.1126/sciadv.aaz1580.

[42] “Epigenetic Control of Smooth Muscle Cell Identity and Lineage Memory | Arteriosclerosis, Thrombosis, and Vascular Biology.” https://www.ahajournals.org/doi/10.1161/ATVBAHA.115.305044 (accessed May 08, 2020).

[43] E. J. Arnoys and J. L. Wang, “Dual localization: Proteins in extracellular and intracellular compartments,” Acta Histochemica, vol. 109, no. 2, pp. 89–110, Apr. 2007, doi: 10.1016/j.acthis.2006.10.002.

[44] R. S. Lim et al., “Multimodal CARS microscopy determination of the impact of diet on macrophage infiltration and lipid accumulation on plaque formation in ApoE-deficient mice,” J Lipid Res, vol. 51, no. 7, pp. 1729–1737, Jul. 2010, doi: 10.1194/jlr.M003616.

[45] B. Feng, D. Zhang, G. Kuriakose, C. M. Devlin, M. Kockx, and I. Tabas, “Niemann-Pick C heterozygosity confers resistance to lesional necrosis and macrophage apoptosis in murine atherosclerosis,” PNAS, vol. 100, no. 18, pp. 10423–10428, Sep. 2003, doi: 10.1073/pnas.1732494100.

[46] Y. Baumer et al., “Ultramorphological analysis of plaque advancement and cholesterol crystal formation in Ldlr knockout mouse atherosclerosis,” Atherosclerosis, vol. 287, pp. 100–111, Aug. 2019, doi: 10.1016/j.atherosclerosis.2019.05.029.

[47] J. D. Hutcheson, N. Maldonado, and E. Aikawa, “Small entities with large impact: microcalcifications and atherosclerotic plaque vulnerability,” Current Opinion in Lipidology, vol. 25, no. 5, pp. 327–332, Oct. 2014, doi: 10.1097/MOL.0000000000000105.

[48] M. Rattazzi et al., “Calcification of Advanced Atherosclerotic Lesions in the Innominate Arteries of ApoE-Deficient Mice,” Arteriosclerosis, Thrombosis, and Vascular Biology, Jul. 2005, doi: 10.1161/01.ATV.0000166600.58468.1b.

[49] M. C. Blaser and E. Aikawa, “Roles and Regulation of Extracellular Vesicles in Cardiovascular Mineral Metabolism,” Front. Cardiovasc. Med., vol. 0, 2018, doi: 10.3389/fcvm.2018.00187.

[50] D. Wolf et al., “Pathogenic Autoimmunity in Atherosclerosis Evolves From Initially Protective Apolipoprotein B100–Reactive CD4+ T-Regulatory Cells,” Circulation, vol. 142, no. 13, pp. 1279–1293, Sep. 2020, doi: 10.1161/CIRCULATIONAHA.119.042863.

[51] S. Hellberg et al., “Alternate models of acute dyslipidemia reveal divergent pathways upon atherosclerosis initiation,” bioRxiv, p. 2020.09.02.279596, Sep. 2020, doi: 10.1101/2020.09.02.279596.

[52] J. Albarrán-Juárez, H. Kaur, M. Grimm, S. Offermanns, and N. Wettschureck, “Lineage tracing of cells involved in atherosclerosis,” Atherosclerosis, vol. 251, pp. 445–453, Aug. 2016, doi: 10.1016/j.atherosclerosis.2016.06.012.

[53] K. R. Stenmark et al., “The adventitia: essential regulator of vascular wall structure and function,” Annu. Rev. Physiol., vol. 75, pp. 23–47, 2013, doi: 10.1146/annurev-physiol-030212-183802.

[54] D. G. Sedding et al., “Vasa Vasorum Angiogenesis: Key Player in the Initiation and Progression of Atherosclerosis and Potential Target for the Treatment of Cardiovascular Disease,” Front Immunol, vol. 9, p. 706, 2018, doi: 10.3389/fimmu.2018.00706.

[55] J. F. Bentzon and M. W. Majesky, “Lineage tracking of origin and fate of smooth muscle cells in atherosclerosis,” Cardiovasc. Res., vol. 114, no. 4, pp. 492–500, 15 2018, doi: 10.1093/cvr/cvx251.

[56] D. Hu et al., “Artery Tertiary Lymphoid Organs Control Aorta Immunity and Protect against Atherosclerosis via Vascular Smooth Muscle Cell Lymphotoxin β Receptors,” Immunity, vol. 42, no. 6, pp. 1100–1115, Jun. 2015, doi: 10.1016/j.immuni.2015.05.015.

[57] J. C. Graver, A. M. H. Boots, E. A. Haacke, A. Diepstra, E. Brouwer, and M. Sandovici, “Massive B-Cell Infiltration and Organization Into Artery Tertiary Lymphoid Organs in the Aorta of Large Vessel Giant Cell Arteritis,” Front. Immunol., vol. 0, 2019, doi: 10.3389/fimmu.2019.00083.

[58] D. Hu, C. Yin, S. Luo, A. J. R. Habenicht, and S. K. Mohanta, “Vascular Smooth Muscle Cells Contribute to Atherosclerosis Immunity,” Front Immunol, vol. 10, p. 1101, 2019, doi: 10.3389/fimmu.2019.01101.

[59] R. A. van Dijk et al., “Systematic Evaluation of the Cellular Innate Immune Response During the Process of Human Atherosclerosis,” J Am Heart Assoc, vol. 5, no. 6, 16 2016, doi: 10.1161/JAHA.115.002860.

[60] S. Y. Park et al., “Resistin derived from diabetic perivascular adipose tissue up-regulates vascular expression of osteopontin via the AP-1 signalling pathway,” The Journal of Pathology, vol. 232, no. 1, pp. 87–97, 2014, doi: 10.1002/path.4286.

[61] J. M. Boucher et al., “Pathological Conversion of Mouse Perivascular Adipose Tissue by Notch Activation,” Arteriosclerosis, Thrombosis, and Vascular Biology, vol. 40, no. 9, pp. 2227–2243, Sep. 2020, doi: 10.1161/ATVBAHA.120.314731.

[62] J. Lin et al., “A Role of RIP3-Mediated Macrophage Necrosis in Atherosclerosis Development,” Cell Reports, vol. 3, no. 1, pp. 200–210, Jan. 2013, doi: 10.1016/j.celrep.2012.12.012.

[63] M. Little, S. M. Kipriyanov, F. L. Gall, and G. Moldenhauer, “Of mice and men: hybridoma and recombinant antibodies,” Immunology Today, vol. 21, no. 8, pp. 364–370, Aug. 2000, doi: 10.1016/S0167-5699(00)01668-6.

[64] D. L. Tribble et al., “Increased low density lipoprotein degradation in aorta of irradiated mice is inhibited by preenrichment of low density lipoprotein with alpha-tocopherol,” J Lipid Res, vol. 41, no. 10, pp. 1666–1672, Oct. 2000.

[65] M. Mehrabian, J. H. Qiao, R. Hyman, D. Ruddle, C. Laughton, and A. J. Lusis, “Influence of the apoA-II gene locus on HDL levels and fatty streak development in mice.,” Arteriosclerosis and Thrombosis: A Journal of Vascular Biology, vol. 13, no. 1, pp. 1–10, Jan. 1993, doi: 10.1161/01.ATV.13.1.1.

