## Supplementary figures and images for "SAMD1 Distribution Patterns in Mouse Atherosclerosis Models Suggest Roles in LDL Retention, Antigen Presentation, and Cell Phenotype Modulation"

### Supplemental Figure 1

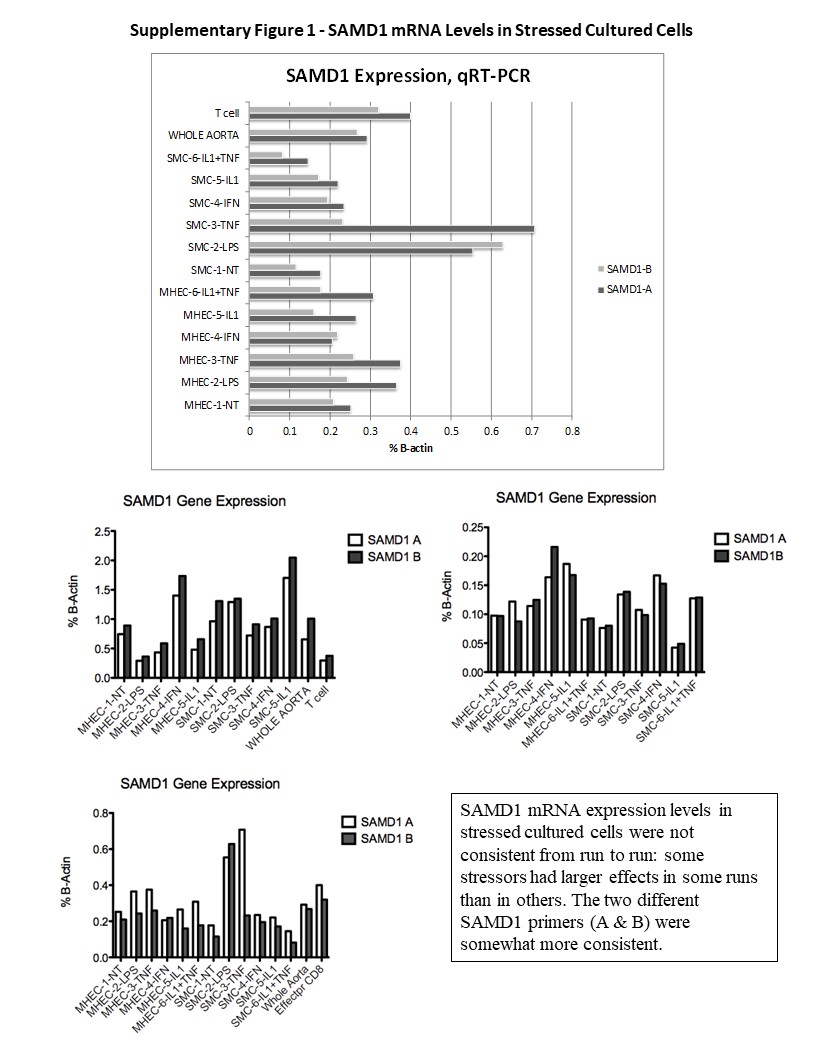
